# Chromosome-level Genomes Reveal the Genetic Basis of Descending Dysploidy and Sex Determination in *Morus* Plants

**DOI:** 10.1101/2022.05.03.490406

**Authors:** Zhongqiang Xia, Xuelei Dai, Wei Fan, Changying Liu, Meirong Zhang, Peipei Bian, Yuping Zhou, Liang Li, Baozhong Zhu, Shuman Liu, Zhengang Li, Xiling Wang, Maode Yu, Zhonghuai Xiang, Yu Jiang, Aichun Zhao

**Affiliations:** State Key Laboratory of Silkworm Genome Biology, Institute of Sericulture and Systems Biology, Southwest University, Chongqing 400716, China; Key Laboratory of Animal Genetics, Breeding and Reproduction of Shaanxi Province, College of Animal Science and Technology, Northwest A&F University, Yangling 712100, China; Key Laboratory of Coarse Cereal Processing, Ministry of Agriculture and Rural Affairs, Chengdu University, Chengdu 610106, China; The Sericultural and Apicultural Research Institute, Yunnan Academy of Agricultural Sciences, Mengzi 661100, China; College of Sericulture, Textile and Biomass Sciences, Southwest University, Chongqing 400716, China

**Author notes:** Equal contributions (Zhongqiang Xia, Xuelei Dai, Wei Fan). Corresponding authors. (Zhao A), (Jiang Y).

**Keywords:** Mulberry, Karyotype evolution, Dioecy, Sex determination, Population genomics

## Abstract

Multiple plant lineages have independently evolved sex chromosomes and variable karyotypes to maintain their sessile lifestyles through constant biological innovation. *Morus notabilis*, a dioecious mulberry species, has the fewest chromosomes among *Morus* spp., but the genetic basis of sex determination and karyotype evolution in this species have not been identified. Three high-quality genome assemblies generated of *Morus* spp. (including those of dioecious *M. notabilis* and *Morus yunnanensis*) were within the range 301-329 Mb in size which were grouped into six pseudochromosomes. Using a combination of genomic approaches, we showed that the putative ancestral karyotype of *Morus* was close to 14 protochromosomes, and that several chromosome fusion events resulted in descending dysploidy (2n = 2x = 12). We also characterized a ∼6.2-Mb sex-determining region on chromosome 3. The four potential male-specific genes, including a partially duplicated DNA helicase gene orthologue (named *MSDH*) and three *Ty3_Gypsy* long terminal repeat retrotransposons (named *MSTG*), were solely identified in the Y-linked area and considered to be strong candidate genes for sex determination or differentiation. Population genomic analysis showed that Guangdong accessions in China were genetically similar to Japanese accessions of mulberry. In addition, genomic areas containing selective sweeps that distinguish domesticated mulberry trees from wild populations in terms of flowering and disease resistance were identified. Our findings provide an important genetic resource for sex identification and molecular breeding in mulberry.

## Introduction

Mulberry (*Morus* spp.) is a major member of the family Moraceae, which also includes various other important plant species, such as banyan tree and paper mulberry. One of the earliest domesticated plants, mulberry, is considered an “Oriental sacred wood” because it is not only a good food source for rearing silkworms but has also been utilized in a number of other ways, including as a fruit, for landscaping, in medicine, in ecological protection, and as forage for animal production [1–3]. The draft genome sequence assembly of male *Morus notabilis* was published in 2013 based on short sequencing reads [4]. In 2020, genome sequencing research revealed the presence of two diploid karyotypes in mulberry species [5]. Congeneric species commonly display varying chromosome counts due to two opposing trends: 1) an increased chromosome copy number as a result of polyploidy (whole-genome duplication [WGD]) and 2) a reduced basic chromosome number *via* structural rearrangements (descending dysploidy) [6, 7]. These processes have long been considered important in speciation due to the shock of chromosome number discrepancies in reproductive isolation [8]. Multi-karyotype evolutionary models for mulberry are limited due to the existence of only few comparative analyses of multiple chromosome-scale genomes. Without considering genomic synteny and large-scale rearrangement events, misleading conclusions on pathway evolution have likely been obtained from evolutionary and comparative analyses of genes for important agronomic traits. Similar to *M. notabilis*, *Morus yunnanensis* is also commonly found in Southwest China due to its unique altitude and humidity requirements [9]. The phylogenetic relationship between wild *M. notabilis* and *M. yunnanensis* is unknown. In addition, our previous study revealed the population structure of cultivated mulberry but lacked the sampling of gene pools within wild mulberry and in the Guangdong region, leaving gaps in our knowledge and raising important questions about the evolutionary history of domesticated perennial mulberry [5].

Mulberry species are either dioecious or hermaphroditic, thus providing abundant resources for research into plant sex determination [10, 11]. The sex determination system of mulberry is similar to that in humans (XY type) [12]. Specific DNA markers associated with the sex determination of male flowers in mulberry have been identified using restriction site-associated DNA (RAD) sequencing [13]. However, the genotypes of these identified markers were not correlated with sex determination among cultivars, and genomic evidence remains unavailable. Previously, clonal propagation was widely used in mulberry propagation, and consequently, sex determination has been little studied in this genus. Moreover, the key genetic basis of sex chromosomes in mulberry has yet to be discovered. With the development of the fruit mulberry industry, cultivation of mulberry cultivars with stable sex characteristics has become increasingly important to ensure high fruit production. Knowledge of sex-specific gene expression is a prerequisite for developing an effective breeding programme for mulberry (especially fruit mulberry). Dissecting the genetic mechanisms underlying dioecy (i.e., separate male and female trees) is crucial for understanding the evolution of this widespread reproductive strategy. *M. notabilis* always show dioecious, which is not affected by the environment, making it an ideal system for studying the genetic basis of sex determination and sex chromosome evolution.

In this study, we performed *de novo* assembly of two chromosome-level genomes of *M. yunnanensis* and female *M. notabilis* to improve the previous draft sequence of male *M. notabilis*, and obtain insights into the evolutionary genomic basis of loci controlling descending dysploidy and sex determination. Compared with our previously reported *M. alba* genome, the two wild species genomes were more diverse in terms of chromosome number and levels of environmental adaptation, providing new insights into karyotype evolution in *Morus*. Through a multidata combination analysis, we not only identified the putative sex-determining region (SDR) and candidate loci responsible for sex determination but also provided a clear framework for broader future studies of sex determination in mulberry species. Population genomic analyses further revealed the phylogenetic relationships within the gene pool of different *Morus* species and the genetic architecture of domestication traits of cultivated mulberry. These sequencing results will provide a valuable resource for future mulberry research and breeding programs.

## Results

### Assembly and annotation of three high-quality *Morus* genomes

To assemble the genomes of male and female individuals of dioecious *M. notabilis* and female *M. yunnanensis*, we used a combination of sequencing technologies including Pacific Biosciences (PacBio) long reads, Illumina paired-end reads, and high-throughput chromosome conformation capture (Hi-C) sequencing reads. Based on k-mer counting, the estimated genome sizes of *M. notabilis* and *M. yunnanensis* were approximately 280 and 296 Mb, respectively (Figure S1A–C). The genome assemblies of female *M. notabilis* and *M. yunnanensis* were highly contiguous, with 96.4% and 95.1% of genome contigs anchored to chromosomes by Hi-C scaffolding, respectively (with genome assembly sizes of 301 and 313 Mb and contig N50 values of 2.7 and 6.5 Mb, respectively). Moreover, Hi-C scaffolding revealed that 91.8% of the original male *M. notabilis* sequences were anchored on six pseudochromosomes with a contig N50 of 6.8 Mb, indicating considerably improved assembly contiguity compared with the original version (Figure S1D–F, Table S1). The coverage of the genome assembly obtained here was evaluated using high-coverage Illumina sequencing data and transcriptome reads mapped against the assembled genome (Table S2). The scores of the long terminal repeat-retrotransposon (LTR-RT) assembly index [14] for male and female *M. notabilis*, and *M. yunnanensis* were 19.98, 20.48, and 21.25, respectively, and the Benchmarking Universal Single-Copy Orthologs (BUSCO) values were 93.5%, 93.6%, and 94.1%, respectively. This result implied that the three genomes were of high quality (**Table 1**).

**Table 1.** Assembly and annotation of the *M. notabilis* and *M. yunnanensis* genomes compared with previously reported genome assemblies.

|  | *M. notabilis*<br>(female) | *M. notabilis*<br>(male) | *M. yunnanensis*<br>(female) | *M. alba*<br>[5] | *M. notabilis*<br>[4] |
| --- | --- | --- | --- | --- | --- |
| **Assembly** |  |  |  |  |  |
| Genome size (bp) | 301,544,460 | 329,129,568 | 313,175,542 | 346,393,484 | 320,378,613 |
| Number of contigs | 539 | 157 | 234 | 398 | 46,842 |
| N50 of contigs (bp) | 2,710,835 | 6,854,161 | 6,538,880 | 2,710,056 | 40,438 |
| GC content of the genome (%) | 0.3511 | 0.3598 | 0.3558 | 0.3429 | 0.3486 |
| Complete BUSCOs | 93.6% | 93.5% | 94.1% | 94.3% | 92.2% |
| Chromosome-scale scaffolds (bp) | 290,812,449 | 302,180,740 | 298,102,870 | 326,128,411 | NA |
| Anchor rate | 96.4% | 91.8% | 95.1% | 94.1% | NA |
| LAI | 20.48 | 19.98 | 21.25 | 18.58 | 8.09 |
| **Annotation** |  |  |  |  |  |
| Number of predicted genes | 25,391 | 25,333 | 2,4851 | 22,767 | 29,338 |
| Average gene length (bp) | 3,140.73 | 3,150.88 | 3,121.06 | 3,209 | NA |
| Average CDS length (bp) | 1,164.39 | 1,160.71 | 1,172.80 | 1,148 | NA |
| Average exon number | 4.86 | 4.83 | 5.00 | 5.09 | NA |
| Complete BUSCOs | 96.1% | 95.6% | 96.0% | NA | NA |
| Repeat sequences (bp) | 164,410,597 | 192,806,284 | 166,442,299 | 180.11 Mb | 128 Mb |
| Annotated to Swiss-Prot | 21,959 | 24,142 | 23,820 | 17,381 | 17,826 |
| Annotated to InterPro | 24,275 | 24,121 | 23,804 | 17,551 | 17,681 |
| Annotated to NR | 14,318 | 24,139 | 23,818 | 22,109 | 22,458 |
| Annotated to TrEMBL | 24,293 | 24,154 | 23,827 | 22,109 | NA |
| Annotated to KEGG | 24,258 | 24,109 | 23,752 | 13,124 | 12,688 |

*Note:* The assemblies of dioecious *M. notabilis* and *M. yunnanensis* were compared with two previously reported genome assemblies of *M. alba* and *M. notabilis* (contig). NA indicates that data were not included in the original articles

For genome annotation, repetitive sequences in the genome were initially annotated by combining *de novo* and homology-based predictions (Table S3). Protein-coding genes were further annotated by combining *ab initio*, homology, and transcriptome analysis methods (Table S3). In total, 25,333, 25,391, and 2,4851 protein-coding genes had 95.6%, 96.1%, and 96.0% BUSCO completeness in male *M. notabilis*, female *M. notabilis*, and female *M. yunnanensis*, respectively, with a total of 98.2%, 98.2% and 98.8%, respectively, of these genes then being functionally annotated in public databases (Table 1).

### Repetitive elements drive genome expansion of mulberry

Syntenic analysis of two *Morus* genomes with grape (*Vitis vinifera*) did not reveal whole-genome duplication (WGD) after the triplication event shared by eudicots, similar to the pattern previously observed in the *M. alba* genome [5] (Figure S2). Furthermore, the collinearities of the genomes of *M. notabilis*, *M. yunnanensis,* and *Ficus benghalensis* (banyan, another member of Moraceae) revealed a high frequency of chromosomal rearrangement events, confirming that *M. notabilis* and *M. yunnanensis* were diploid (2n = 2x = 12) (**Figure 1A** and 1B). Phylogenomic analysis of eight angiosperms using single-copy orthologues identified by OrthoFinder [15] revealed that *M. notabilis* and *M. yunnanensis* diverged only approximately 3 million years ago (Mya) (Figure S3), indicating their close phylogenetic relationship. Combined with similar histological morphologies (Figure S4), these results suggest that they belonged to the same subgenus. Moreover, using a combination of several wild mulberry accessions and *Ficus* as the outgroup, we constructed an evolutionary tree with single nucleotide polymorphisms (SNPs) of four-fold degenerate (neutral) evolving sites based on the maximum likelihood method in IQ-TREE [16]. The evolutionary distances and divergence times between *M. notabilis* and domesticated *M. alba* were far greater than those observed among other wild mulberry trees (Figure 1C), and these two species may belong to parallel evolutionary clades.

**Figure 1.**
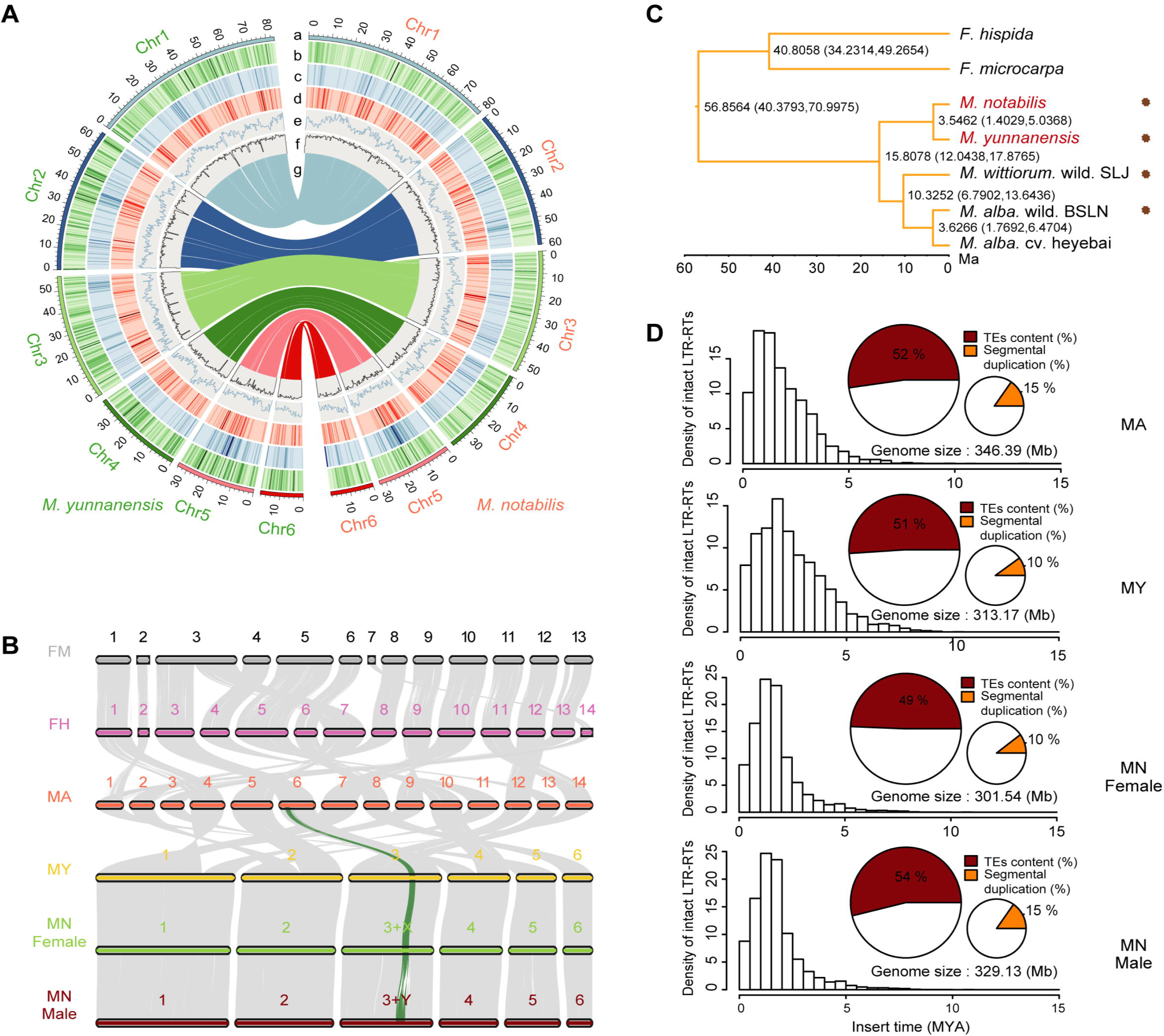
Genome evolution in the genus *Morus.* **A.** Genome landscape of *M. notabilis* and *M. yunnanensis*. The tracks indicate parameters described below. a. Chromosome karyotypes of *M. notabilis* and *M. yunnanensis*. The outer circle represents the chromosome length, with units of Mb. b. Segmental duplications. c. Density of all *Gypsy* LTR-RTs. d. Density of all *Copia* LTR-RTs. e. Gene density. f. GC content. g. Synteny between the two genomes. **B**. Chromosome synteny between *F. microcarpa* (FM), *F. hispida* (FH), *M. alba* (MA), *M. yunnanensis* (MA), and *M. notabilis* (MN), with chromosome names shown above. The green line indicates the synteny blocks in the Y-linked regions of female and male *Morus*. **C.** Phylogenetic relationship among different subgenera of mulberry. A maximum-likelihood tree was constructed by extracting neutrally evolving sites from resequenced data from *M. notabilis* C.K. Schneid (CS)*, M. yunnanensis* Koidz*, M. wittiorum* L. wild. SLJ*, M. alba* L. wild. BSLN, and *M. alba* L. cv. Heyebai (*F. microcarpa* and *F. hispida* as outgroups). The divergence times among different groups of species are labelled on the right. Clade support values near nodes represent the estimates of divergence time (Mya) with a 95% credibility interval. The asterisk represents the wild species. **D.** LTR burst patterns and fractions of TEs and SD in *M. alba*, *M. yunnanensis*, and female and male *M. notabilis*.

The whole-genome alignment analysis showed that 30.2 Mb (9% of the assembled genome) covering 870 genes in *M. notabilis* and 51.4 Mb (15%) covering 948 genes in *M. alba* were segmental duplications (SDs) (Figure 1D and Table S3), indicating that SD was a major contributor to genome-size expansion. Genes overlapping within the female *M. notabilis* SD regions were significantly enriched in fatty acid degradation biological processes (“ath00071”, hypergeometric test, corrected *P value* < 8.10×10^−5^) (Figure S5). We identified 58.45% of the male *M. notabilis* genome, 54.16% of the female *M. notabilis* genome, and 56.93% of the *M. yunnanensis* genome as transposable elements (TEs) (Table S3). Genome-wide proliferation of intact LTR-RTs from *M. yunnanensis* and female and male *M. notabilis* occurred approximately 1.5 Mya. Moreover, these LTR-RTs were dated more recently than *M. alba* formation (Figure 1D and Table S3). Therefore, we concluded that a high level of divergence in SDs and TEs existed between *M. notabilis* and *M. alba*.

We also identified an average of 13,179,790 SNPs between the *M. notabilis* and *M. alba* genomes in one-to-one aligned regions (Table S4). The number of small insertions and deletions (50–500 bp) in those one-to-one aligned regions was 1,479,461 bp on average, which accounted for approximately 0.49% of the *M. notabilis* genome (Table S4). Notably, fewer SNPs (5,186,831), small insertions, and insertion–deletion mutations (indels) were observed between the *M. notabilis* and *M. yunnanensis* genomes than between the *M. notabilis* and *M. alba* genomes, suggesting a strong relationship between the two wild-grown plants, a finding consistent with the results from the phylogenetic tree described above. Two sequences of 31.91 Mb and 125.95 Mb were affected by structural variants (SVs) in the two comparisons. In particular, the majority (71%) of the SV sequences in the comparison between *M. notabilis* and *M. alba* showed repeat expansion and contraction.

### Chromosomal fusion in mulberry genomes is associated with adaptive evolution

Chromosomal evolution is associated with genome size, gene family evolution, and speciation. Genome structural changes led to the present-day karyotypes of *M. notabilis* (2n = 12) and *M. alba* (2n = 28). Using the available genomes of *M. notabilis*, *M. alba*, *F. microcarpa*, *F. hispida*, and *Cannabis sativa* (as the outgroup), we reconstructed the ancestral karyotype of the Moraceae (n = 21) (**Figure 2A**), which corresponded to the ancestral eudicot karyotype (AEK) [17]. Compared with the putative ancestral chromosomes, 39 and 40 large syntenic blocks were identified in *M. notabilis* and *M. alba*, respectively (Table S5), which enabled us to deduce the arrangements of ancestral chromosome segments in mulberry. Karyotyping of *M. alba* revealed that at least ten major chromosomal fusions (CFUs) and one chromosomal fission of 21 chromosomes of the palaeohexaploid ancestor may have been involved. Approximately half of the chromosomes of *M. alba* were found to have descended from a single ancient chromosome, a finding similar to the results reported for paper mulberry [18]. Moreover, pseudochromosomes of *M. notabilis* were constructed from the ancestor karyotype *via* at least 18 CFU events and one chromosomal fission event, resulting in a substantial decrease in the haploid chromosome number from 21 to six. Furthermore, *Mn*Chr1, *Mn*Chr2, *Mn*Chr3, and *Mn*Chr4 were each derived from at least two *M. alba* chromosomes *via* complex translocations. The high-quality genome assemblies revealed that the karyotype of *M. alba* is more similar than that of *M. notabilis* to the ancestral karyotype, whereas the karyotype of *M. notabilis* was derived from more CFUs. We performed a repeat element annotation analysis to investigate the genome features of evolutionary fusion regions (EFRs) and found that the EFRs in the *M. notabilis* genomes were mainly enriched with LTR elements (50.29%) (Table S6).

**Figure 2.**
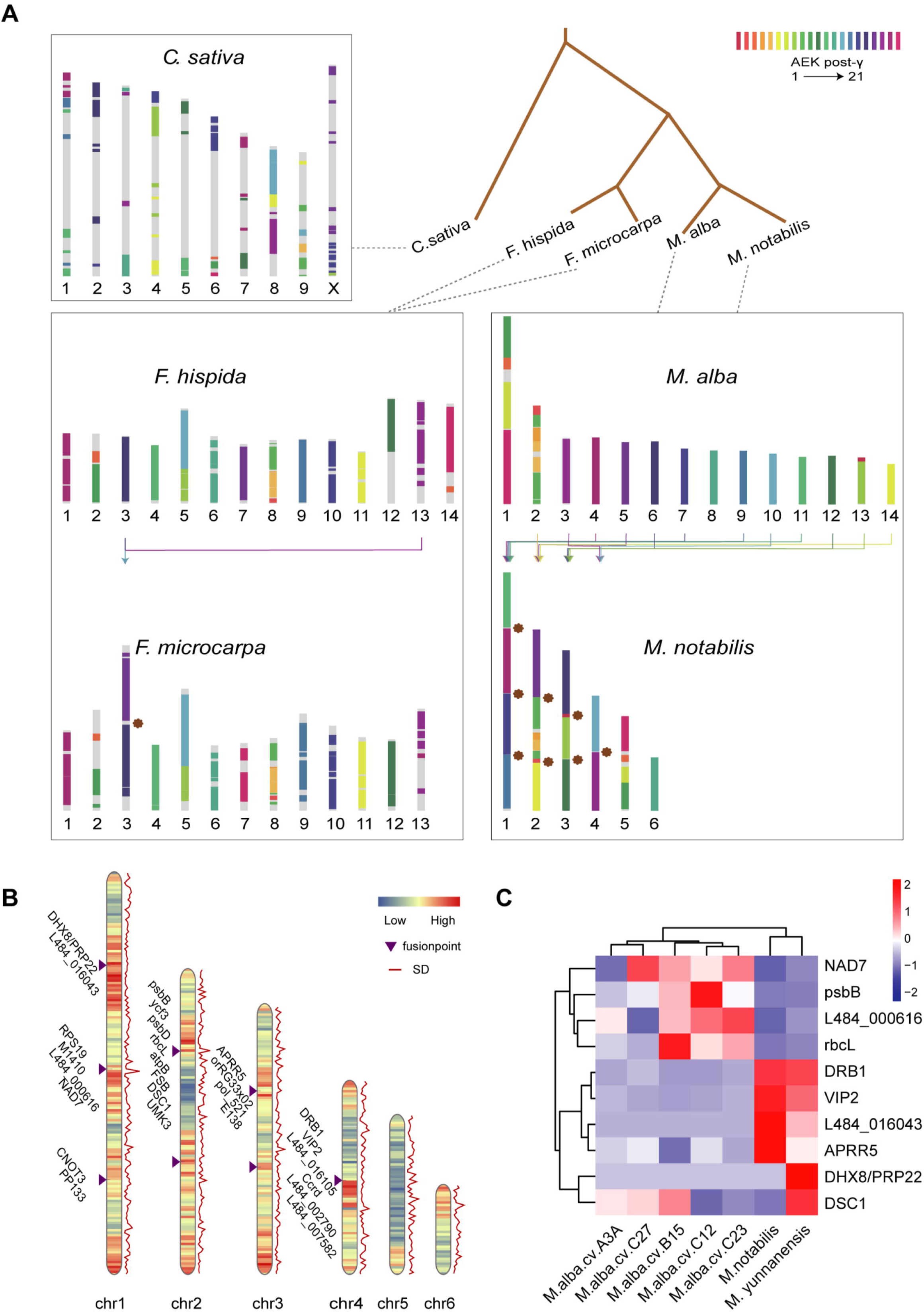
Reconstruction of ancestral chromosomes of Moraceae with *Cannabis sativa* as an outgroup. **A**. Probable distribution of ancestral chromosome segments in the banyan tree and *Morus* genomes according to the ancestral eudicot karyotype (AEK) model proposed by Murat et al. [17]. Blocks are “painted” with colours corresponding to ancestral chromosomes (AEK1–AEK21). Brown asterisks in the *M. notabilis* chromosome diagram indicate sites of chromosomal rearrangement**. B**. Genome-wide landscape of chromosome fusion features in *M. notabilis*. The purple triangle represents the rearrangement site. The heatmap and the red line represent the distribution of gene density and SDs on the chromosomes, respectively. **C**. Transcriptome expression levels of genes in the fusion region of two karyotypes of *Morus* plants.

We identified 26 genes located in six chromosomal fusion regions (Figure 2B). The Kyoto Encyclopedia of Genes and Genomes (KEGG) enrichment analysis confirmed that the genes in the rearranged regions of *M. notabilis* and *M. alba* chromosomes were mainly enriched in functions related to photosynthesis (“ath00195”, hypergeometric test, corrected *P value* < 6.35×10^−10^) and metabolic pathways (“ath01100”, hypergeometric test, corrected *P value* < 1.55×10^−4^) (Figure S6A). The Gene Ontology (GO) annotation results showed that the rearranged regions were mainly involved in adenosine diphosphate-binding sites (“GO: 0043531”, hypergeometric test, corrected *P* value < 4.34×10^−9^) and positive regulation of gene expression (“GO: 0010628”, hypergeometric test, corrected *P* value < 2.13×10^−5^) (Figure S6B), which prompted us to investigate the function of chromosomal shuffling loci in *Morus* species. We then examined the transcriptome expression levels of these genes in *M. notabilis* and the *M. alba* cultivars (Figure 2C and Table S7). Most of these fusion genes were differentially expressed in the two karyotypes of mulberry. These findings provide novel insights into chromosome evolution and link chromosomal rearrangements to the evolution of functional genes.

### Location of sex-determining regions and identification of candidate sex-determining genes

Short Illumina reads of four male and four female plants of *M. notabilis* were subsequently catalogued into 40-bp k-mers in different categories using the genome-wide classification method to identify the putative sex chromosome and sex locus regions [19, 20]. Sex-specific k-mers (detected in all samples of one sex, but not in the other sex) were obtained, including 333,348 male-specific k-mers (MSKs) and 1,664 female-specific k-mers (FSKs). The higher MSK count was consistent with the results obtained for persimmon and ginkgo [19, 20], suggesting unique genomic regions in male individuals. This result indicated that the sex determination system of *M. notabilis* may be an XY system.

Based on the position of male-specific reads in the genome assembly, we identified a candidate sex-determining region (SDR) (Chr3: 38,911,287-45,186,478) that contained male-specific reads with 100-kb windows, whereas female-specific reads were found to be uniformly distributed throughout the genome (**Figure 3Aa**, Figure S7). We then detected higher densities of SNPs and indels in the SDR in male *M. notabilis* individuals than in female individuals but not in the rest of the genome, indicating early divergence between Y and X chromosomes (Figure 3Ac-e). In addition, we further analysed the genome-wide methylation levels of male and female flowers, and found that CG and CHG methylation levels were higher in the candidate SDR of males (Figure 3Af-h).

**Figure 3.**
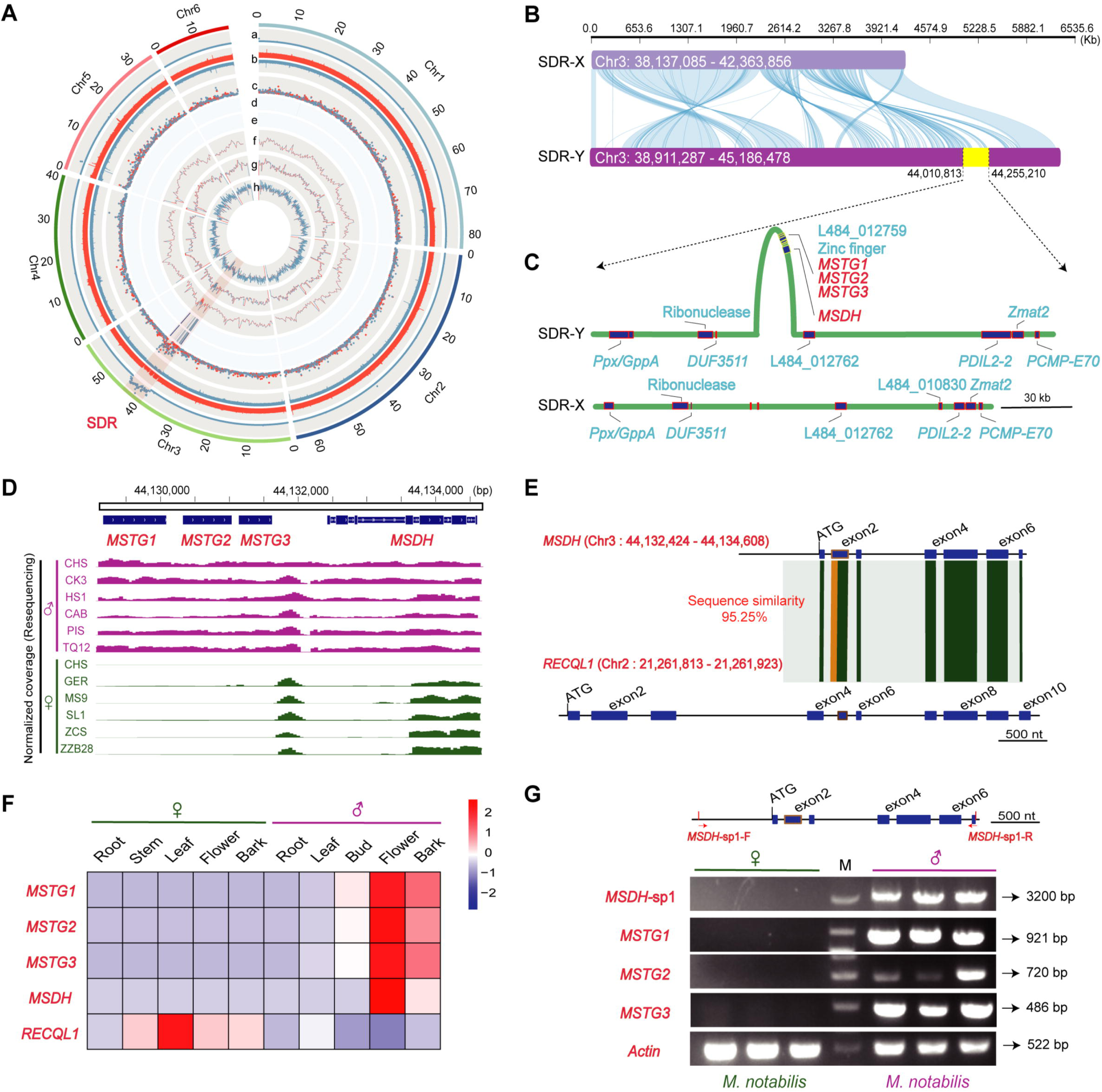
Reconstructed haplotypes in the SDR and identification of candidate sex-determining genes in *M. notabilis.* **A.** Identification of the sex chromosome and SDR among six chromosomes of the *M. notabilis* genome. The tracks indicate the parameters described below. (a) Manhattan plots of the mapping depths of MSKs in the male *M. notabilis* genome (100-kb window). (b) Mapping coverage of female (red) and male (blue) *M. notabilis* Illumina reads. (c) SNP and indel densities in female (red) and male (blue) *M. notabilis*. (d) Heatmap showing the density of candidate male-specific SNPs. (e) Heatmap showing the density of candidate male-specific indels with the same colour coding as track d. (f-h) Whole-genome methylation levels in CG, CHG, and CHH contexts are shown. **B.** SDR-X and SDR-Y haplotypes were reconstructed from genome sequences in our assembly. The plot was created using RectChr (https://github.com/BGI-shenzhen/RectChr). **C.** Gene distribution diagram of candidate regions. **D.** Validation of four candidate sex-specific genes in two sex phenotypes (six females and six males) using IGV. Red represents males, and green represents females. The top panel presents the structure of genes. **E.** Schematic depicting the events leading to the formation of the *M. notabilis MSDH1* gene, which is likely to result from alternative splicing in the *RECQL1* gene. The shaded block indicates the duplicated segments described in the text. **F.** Transcriptome profiles of four sex-specific candidate genes. All genes were expressed at high levels in male flower tissues. **G.** Agarose gel electrophoresis profile of four candidate genes in females and males. The location of the *MSDH*-specific 1 (*MSDH*-sp1) primer is shown at the top. Actin was used as the control. M, molecular marker. The primer sequences are listed in Table S10.

We delineated the SDR-X and SDR-Y haplotypes in this region (Figure 3B). The collinearity results indicated that some fragments of the X and Y haplotypes were not aligned, which motivated us to identify any inversions present on the Y chromosome in *M. notabilis* that masked recombination, consistent with the results obtained from the banyan tree [21]. Based on this result, these sex-specific regions contained additional candidate genes for sex determination. From the analysis, 404 genes were predicted in the SDR-Y haplotype, and differentiation of Y and X haplotypes in this region provided strong evidence for the presence of a fully linked region on Chr3. A total of 306 conserved gene pairs (75.76%) from synteny blocks within the SDR and X counterpart were detected using MCscan (Figure S8 and Table S8). We further evaluated two genomic data sets for sex determination as follows: 1) based on resequencing data of different sexes, we determined the regions present only in male *morus* genomes, and 2) the expression of sex bias-related genes in male flowers was measured. Notably, all analyses revealed the existence of a 5-kb region (Chr3: 44,129,165-44,134,608; Figure 3C and 3D; Figure S9 and S10).

Manual annotation and curation revealed four genes in this region: three male-specific *Ty3_Gypsy* RT gene models (designated *MSTG1, MSTG2,* and *MSTG3*) and one male-specific DNA helicase gene (designated *MSDH*) resulting from partial duplication of the gene *RECQL1* (located on Chr2 and found in both male and female genomes) (Figure 3E). The transcriptomic analysis showed that the four male-specific genes were expressed at high levels in male flowers, suggesting their importance in male development and maintenance in *M. notabilis* (Figure 3F and Table S9).

In particular, the coding region of the *MSDH* gene showed 95% nucleotide identity with that of canonical *RECQL1,* with considerable divergence at the N-terminus. The *RECQL1* gene contained 10 exons, while *MSDH* had only seven exons (Figure 3F). Moreover, the *MSDH* gene in *M. notabilis* appeared to have undergone a splicing event involving its second exon. *MSDH* was predicted to have helicase_ATP_BIND and helicase_C domains and structural similarities to *RECQL1* (Figure S11), which are critical for anti-crossover signalling during meiotic recombination [22].

We used polymerase chain reaction (PCR) primers designed to amplify four separate gene fragments and detected the presence of target fragments of each sex in a collection of dioecious mulberry species, including *M. notabilis* and species from other subgenera. For the *MSDH* gene, we designed a specific gene primer fragment (named *MSDH*-sp1; physical location shown in Figure 3G) for male-specific amplification. Four gene fragments were successfully amplified in all male individuals, whereas no amplified products were obtained from female individuals (Figure 3G and Figures S12–S13). We also surveyed the in-house and publicly available genome assemblies of mulberry species; the results reveal that the four candidate genes were well supported in the male genome but were absent in the female genome (Figure S14). Based on these results, we revealed a possible XX/XY- determining system and putative candidate sex-determining genes in *Mours*.

### Population structure analysis improves the landscape of genetic affinity in mulberry accessions

Wild relatives are expected to be important sources of genetic diversity in mulberry, and understanding the phylogenetic relationship between wild and cultivated mulberry accessions might greatly facilitate mulberry breeding. In the present study, we resequenced 32 representative wild and landrace specimens from various regions, including Cambodia, Sri Lanka, China, and other countries. Using these data and publicly available genomic sequences for 123 cultivars/landraces, the genetic divergence among multiple wild and domesticated species was studied (Table S12). All sequence reads were mapped to the *M. alba* genome with an average coverage depth of ∼21.6× (Table S13), and a total of 29,185,577 SNPs were identified and used in subsequent population-based genomic analyses (Table S14 and Figure 4B).

**Figure 4.**
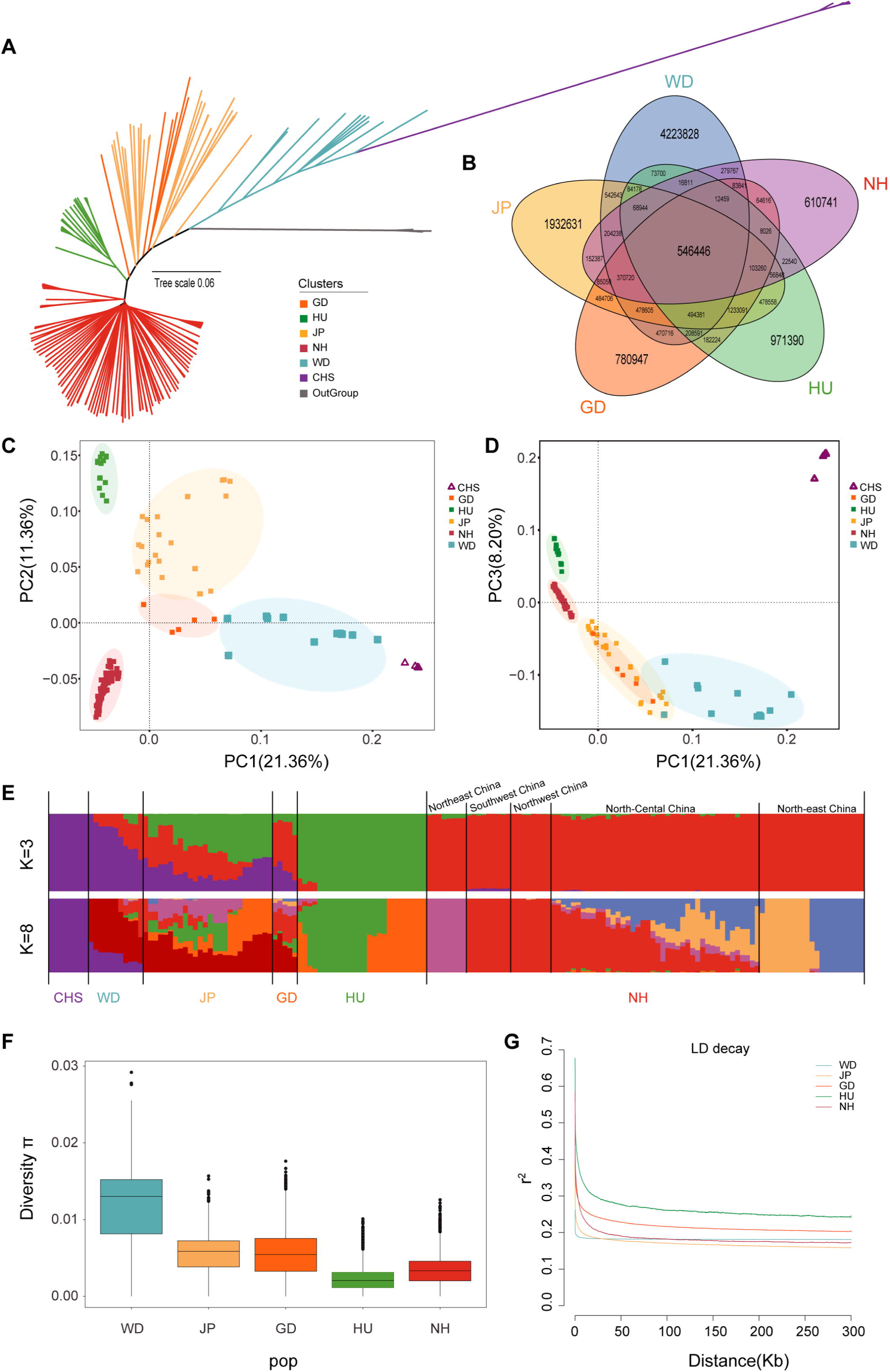
Genetic diversity of mulberry accessions. **A.** NJ tree from the genome sequences used in this study. **B.** Venn diagrams showing the number of SNPs of WD, JP, GD, HU, and NH. Arbitrary colours were used to better visualize the different groups, and the overlapping areas represent SNPs shared between groups. **C** and **D.** PCA of wild and domestic accessions: C (PC1-PC2) and D (PC1-PC3). **E.** Model-based clustering of wild and domestic mulberry trees using ADMIXTURE with k = 3 and k = 8. The colours are the same as those used in panels A, C and D. **F.** Genome-wide distribution of the nucleotide diversity of each group in 50-kb windows with 50- kb steps. The horizontal line inside the box indicates the median of this distribution, the box limits indicate the first and third quartiles, and the points show outliers. **G.** Genome-wide average LD decay estimated for each group.

We performed an ADMIXTURE analysis, neighbour-joining (NJ) tree analysis, and principal component analysis (PCA) using genomic SNPs to investigate the phylogenetic relationships between wild relatives and cultivated mulberry. The NJ tree showed clustering of mulberry accessions into six separate genetic groups with the *Ficus* genus (including *F. hispida* and *F. macrocarpa*) at the root (**Figure 4**A). The first group (CHS) included the wild-growing plants *M. notabilis* and *M. yunnanensis* from Southwest China, the second group (WD) included 11 wild plants collected in China and other countries, and the third group (JP) mainly included landraces and cultivars from Japan and other countries. The fourth group (GD) consisted of landraces and cultivated species mainly from Guangdong Province. The fifth group (HU) was mainly derived from Taihu Basin in the southern Yangtze delta plain in China, while the sixth group (NH) consisted of cultivated mulberry accessions from other places, mainly distributed throughout northern China. The phylogeny of the wild species revealed their close relationships with the cultivated relatives in the JP and GD groups; PCA was performed to confirm these phylogenetic relationships (Figure 4C–D). When higher k values were used (Table S15), the NH group was further divided into subgroups, including those from Northeast China, Southwest China, Northwest China, North-central China, and Northeast China (Figure 4E and Figure S15).

The results presented in Figure 4F show that nucleotide diversity (θπ) was highest in WD, followed by JP and GD (Table S16). The highest θπ (5.51 × 10^−3^) among the four domesticated groups was observed in JP, which also showed the most singletons and the highest linkage disequilibrium (LD) decay rate (Figure 4G). We further explored the phylogeny and migration history of cultivated and wild mulberry using TreeMix [23]. When using CHS as an outgroup, a bifurcation pattern similar to the initial phylogenetic result was observed when up to two migration events were included, and gene flow to HU cultivated mulberry was observed in the JP group (Figure S16).

### Screening for selective sweeps related to domestication

Present-day mulberry cultivars exhibit diversity in many agronomic characteristics, such as flowering time, disease resistance, and leaf development. Among them, traits such as resistance strength and delayed flowering have been recognized as important agronomic traits in cultivated mulberry. We compared domestic (JP, GD, HU, and NH) and wild mulberry (WD) populations based on the fixation index (*F*_ST_) and nucleotide diversity in 50-kb sliding windows of the genome (**Figure 5**A). We defined the windows with outlier signals (top 1%) for both statistics (*F*_ST_ > 0.564, π ln-ratio WD/JP-GD-HU-NH > 2.041) as harbouring putative selective sweeps. Merging outlier windows yielded 103 unique regions that contained 411 positively selected genes. We also performed a functional enrichment analysis by identifying GO terms for these overlapping genes (Table S17). The GO analysis revealed three significantly enriched biological processes (hypergeometric test, corrected *P* value < 0.01) that were associated with disease resistance (corrected *P* value = 3.10×10^−5^ to 2.42×10^−9^), flowering (corrected *P* value = 7.52×10^−6^ to 4.67×10^−7^), and plant hormones (corrected *P* value = 4.41×10^−5^ to 2.02×10^−6^) (Figure 5B).

**Figure 5.**
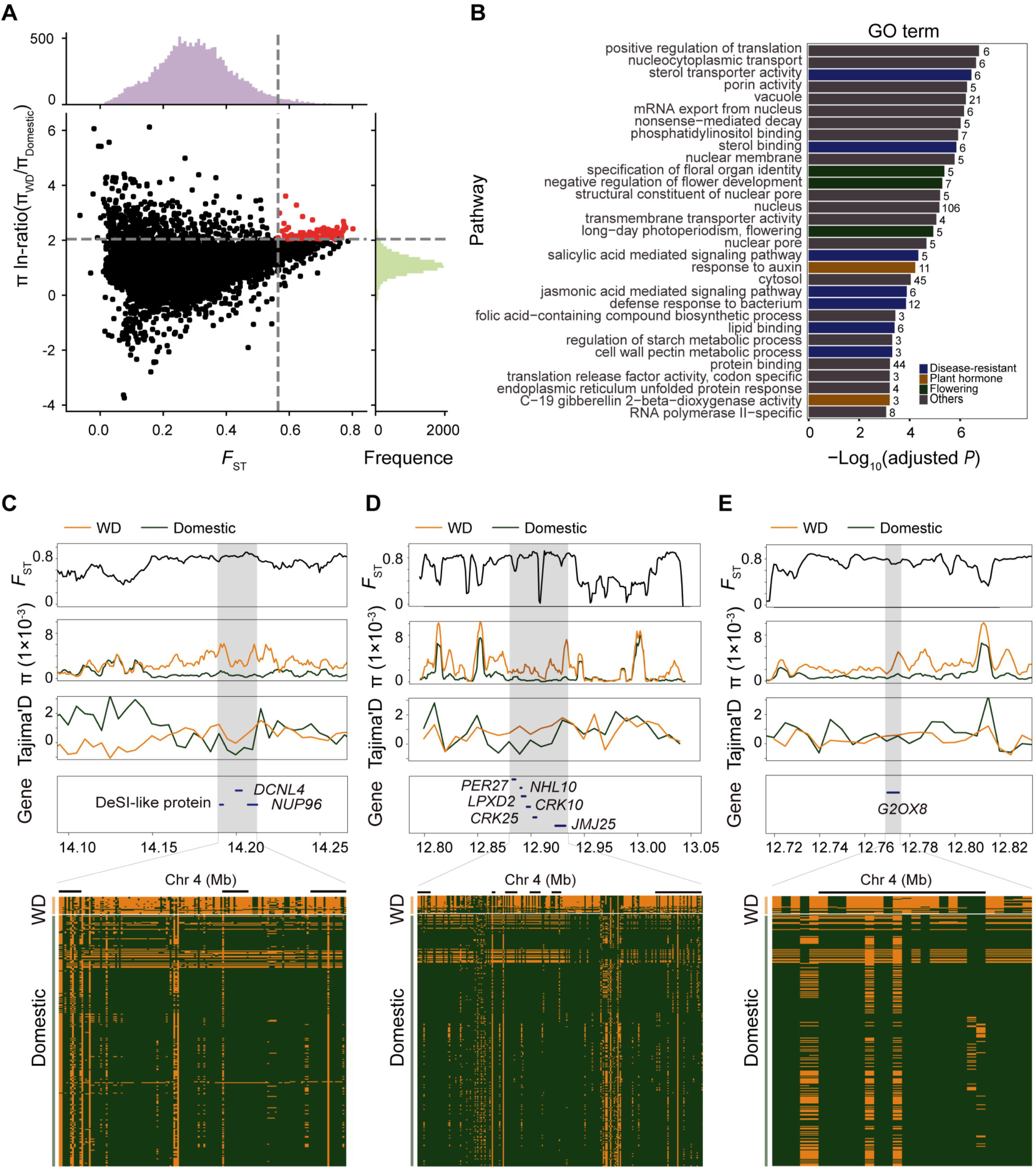
Selection signatures identified from comparisons between wild and domestic mulberry. **A.** Distribution of the pairwise fixation index (*F*_ST_) (x-axis) and π ln ratio (y-axis) between WD and domestic mulberry trees. The dashed vertical and horizontal lines indicate the significance thresholds (*F*_ST_ > 0.564, π ln-ratio WD/JP- GD-HU-NH > 2.041). **B**. GO terms identified as significantly overrepresented (hypergeometric test, corrected *P* value < 0.01). **C, D** and **E.** *F*_ST_, nucleotide diversity and Tajima’s D plots of the three candidate genomic regions. The heatmap on the bottom shows SNPs with minor allele frequencies > 0.05 that were used to infer haplotype patterns. The major allele at each SNP position in WD is coloured green, and the minor allele is coloured yellow.

Changes in flowering time are a major goal of mulberry breeding programmes for the production of cultivars optimally adapted to local environments. Therefore, we identified three overlapping genes on chromosome 4 that were associated with flowering time in cultivated mulberry (*DCNL4*, *NUP96*, and *DESL-like*) using the three selection methods mentioned above (*F*_ST_ = 0.74 and π ln-ratio WD/domestic = 2.133). This analysis indicated significant selection for these genes in response to the domestication of mulberry. The positive selection signals around this region were further confirmed by significantly lower values for Tajima’s D and long haplotype patterns in WD (Figure 5C). Of these three genes, *NUP96* functions as a negative regulator of long day-induced flowering [24], whereas *DCNL4* is proposed to be involved in pollen development and embryogenesis [25]. Current knowledge of these genes suggested that mulberry domestication involved selection of flowering time.

Mulberry production is severely threatened by diseases, which highlights the importance of breeding programs with a focus on disease resistance for crop improvement. Our selective sweep analyses revealed genomic regions and candidate genes associated with disease resistance, and some selected genes were involved in increasing plant defence responses (e.g., *NHL10*, *LPXD2*, *CRK10/25*, and *JMJ25*) (Figure 5D) [26–31]. Signals for positive selection in cultivars compared with WD were also detected for the gene *G2OX8*, which encodes the gibberellin 2-beta-hydroxylase enzyme, resulting in changes in hypocotyl length or plant height via the gibberellin signalling pathway (Figure 5E) [32, 33]. These results should be valuable to future mulberry research and breeding plans and should exert a positive effect on mulberry improvement.

## Discussion

Genome rearrangements, a common genetic process associated with speciation, have long been postulated to be a key phenomenon in the evolution of higher eukaryotes such as land plants. Recently, with the surge in plant genome sequencing projects and the advances bioinformatics tools, more examples of selection-induced plant genome evolution driven by genome rearrangement, such as in rye (*Secale cereale*) [34], *Brassica* crops [35], and *Miscanthus floridulus* [36], have been reported. Our findings on *M. notabilis* and a previous study [21] have verified that the formation and diversification of karyotypes in plants were possible largely attributed to gene gain and reuse via SVs.

Three high-quality mulberry genomes and their distinct, descending dysploidy status enabled us to infer the origins of and steps leading to the current mulberry karyotype. The ancestral karyotype of mulberry was particularly similar to that of modern *M. alba* (2n = 14) and was most similar to the inferred ancestral chromosomal arrangement, a finding that is consistent with the results reported for paper mulberry [18]. Due to adaptation in response to unique environmental changes, karyotype formation in *M. notabilis* involved more chromosomal fusion events than in other plants and caused the observed descending dysploidy (from x = 14 to x = 6), which indicated parallel evolution and selection throughout the genus. Researchers previously hypothesized that this type of chromosomal fusion-fission event may be a common phenomenon in *M. notabilis* during growth and development [37]. Some of the genes present in evolutionary rearrangement regions may be associated with the adaptation and evolution of the species [38], a hypothesis consistent with our findings. Alternative theories for descending dysploidy generation, such as “telomere sequence directivity”, may also be applicable but will require an analysis with further sequencing data and genome analysis. The two unsolved studies on chromosomal behaviour by Barbara McClintock promoted further experiments on decreasing chromosome numbers, suggesting that centromere breaks, inactivation, and fusions, in combination with telomeres, may be common mechanisms of karyotype evolution in plants and may extend to all eukaryotes [39]. However, mechanisms that coregulate plant chromosomal rearrangements, such as descending dysploidy or WGD, at the telomere and centromere levels remain largely unclear and require more research.

Genetically determined dioecy occurs through mutations in two linked genes, one causing male infertility and the other causing female infertility, according to a theoretical concept of the formation of sex chromosomes [40, 41]. Dual-gene models have been discovered in campion (*Silene* sp.), papaya, asparagus, and kiwifruit in recent empirical studies [42–45]. In persimmon, however, a single gene appears to be sufficient for the expression of male features and inhibition of female development [19]. The pattern of sequence differences between males and females observed in the present study suggested the existence of a large nonrecombining region containing genes involved in sex determination and that male sex in mulberry is associated with heterogamy. We hypothesize that the hemizygosity of this Y-specific region is caused by mutations in sex-determining genes, which might also explain why regions containing sex-determining genes do not recombine between the Y and X haploids. In plants, most sex-biased genes are not carried on sex chromosomes, and their expression levels are most likely regulated by an upstream sex-determining gene or genes [46, 47].

As TE insertion is rarely fixed in all individuals within a species, the presence of TE sequences in the same location is unusual [48]. Insertion of a complete sequence over a long evolutionary period is necessary for males to be retained in a specific area. Three *MSTG* transposon sequences are absent in the genomes of *Arabidopsis thaliana* and *Oryza sativa*. However, they exist stably in the mulberry genus (Figure S12), suggesting that their specific functions are conserved. TE insertions have been shown to silence the expression of neighbouring genes, such as those that control sexual forms in monoecious individuals [49]. This finding is surprising and implies that TE insertions have specialized functions in plants, such as improved male function. Although the specific activities of these genes are unknown, *Ty3_Gypsy* in budding yeast and humans select binding sites for essential meiosis transcription factors, linking their transcriptional activity to the meiosis process [50, 51]. *Ty3_Gypsy* LTR-RTs contribute to the development of *Populus deltoides* flowers of male plants by producing long noncoding RNAs (lncRNAs) [52]. Based on a *de novo* repeat library constructed from *M. notabilis* genome sequences (see the Methods section), *MSTG*s are annotated as transposable proteins in the LTR/Gypsy transposon. Consistent with their functions as transposable proteins, transcripts were detected to have the ability to encode proteins using lncRNA-seq and general RNA-Seq (Figure S17), confirming that these transcripts do not produce lncRNAs. These results suggested that these transposon genes may have new regulatory patterns at the level of sex differentiation.

In plants, DNA helicases, such as *RECQ4A* and *RECQ4B* [22], and their interacting partners play a role in meiotic recombination [53]. In *Drosophila*, one *male-specific lethal* (*msl*) gene encodes a protein with sequence similarity to members of a superfamily of RNA and DNA helicases [54]. *MSDH* and *RECQL1* have very similar sequence structures, with alternative splicing occurring in exon 2 of *MSDH* (Figure 3E). The phylogenetic analysis revealed that the *MSDH* gene is a partial duplicate of the *RECQL1* gene in male *M. notabilis* (Figure S18). The transcriptomic analysis showed that canonical *RECQL1* was expressed in both leaves and female flowers, with a higher expression level in female flowers, whereas *MSDH* expression was specific to male flowers (Figure 3F and Figure S19). Expression data from strand-specific lncRNA-Seq and small RNA-Seq revealed that *MSDH* is transcribed into long transcripts that do not generate small interfering RNAs (siRNAs; Figure S17). We also used bisulfite sequencing to analyse methylation levels in the *RECQL1* gene region and discovered no significant sex-biased differences (Figure S20), implying that *MSDH* and *RECQL1* function independently. *MSDH* is therefore either a new gene that evolved *de novo* or a gene that transposed to new locations, followed by partial loss of the duplicated sequences. However, *MSDH* genes may have a very complex regulatory network, which is currently not well characterized, and this network will be the focus of our future research.

Understanding population structure and phylogenetic relationships is very important for the management and utilization of gene pools of germplasm resources. Despite its widespread usage in basic mulberry research, the CHS group is not of value in mulberry breeding, probably due to reproductive obstacles. Our evolutionary study answered various questions about the taxonomic status of these distantly related species and highlighted a potential future path for germplasm collection and exploitation. Based on the samples surveyed, Japan and the Guangdong region of China were the most promising sources of germplasm resources because the populations in these regions had the greatest nucleotide diversity and shared the same genetic composition, extending our previous understanding of their classification. East Asia has been identified as a significant ancient hotspot for the domestication of crops, including rice, sorghum, millet, soybeans, foxnut, apricot, and peach [55, 56]. The domestication and diversification of mulberry, similar to those of other woody plant species, involve several complex steps, leading to geographic radiation and deliberate breeding of varieties. The selection of traits to maximize yield and quality necessitates the collection of accessions and additional evidence in the future to test our proposed evolutionary scenario [57].

## Materials and methods

### Plant materials

Young leaves of two wild mulberry species, *M. notabilis* (including males and females) and *M. yunnanensis*, were collected for whole-genome and *de novo* assembly. *M. notabilis* grows in Yingjing County, Sichuan Province, China (N 29°80’ E 102°85′, altitude 1100–1400 m), and *M. yunnanensis* grows in Pingbian County, Yunnan Province, China (N 22°68′ E 103°67′, altitude 1,900–2,200 m). Young leaves were collected from wild mulberry species in various regions in China and other countries, grafted, and stored at the Mulberry Breeding Center, Southwest University, for genome resequencing.

### Genome size estimation

Short Illumina reads were obtained to estimate the size of the two genomes using a publicly available PERL script (https://github.com/josephryan/estimate_genome_size.pl) to determine the distribution of k-mer values with Jellyfish [58]. Genome size was calculated by dividing the total number of k-mers by the k-mer distribution peak. For visualization, the online web software GenomeScope [59] was used.

### Genome sequencing and assembly

#### Illumina short-read sequencing

Cetyltrimethylammonium bromide (CTAB) was used to extract genomic DNA [60]. The Illumina HiSeq platform was used to sequence a library with a 350-bp insert size, yielding 150-bp paired-end reads (Table S1). The raw reads were subsequently trimmed and filtered to acquire clean reads.

#### PacBio library construction and sequencing

A portion of the DNA samples was transferred to AnnoRoad (Ningbo, China) for the construction of circular consensus sequence (CCS) libraries (male *M. notabilis*) and 20K *de novo* libraries (female *M. notabilis* and *M. yunnanensis*) using Pacific Biosciences methodology and sequenced using a PacBio Sequel platform.

#### Hi-C library construction and sequencing

The library for Hi-C sequencing was created from young leaves crosslinked with the *Mbo*I restriction enzyme, as described previously [5]. Then, the Hi-C libraries were amplified using 12–14 cycles of PCR and sequenced on the Illumina HiSeq platform. An Illumina HiSeq instrument combined with 2150-bp reads was used to infer the sequencing interaction pattern.

#### Genome assembly and pseudomolecule construction

The genomes mentioned here were assembled as described below. (1) For male *M. notabilis*, we used hifiasm [61] with the default parameters to construct contigs from PacBio HiFi CCS clean reads. PacBio SMRT subreads were corrected and assembled into contigs for the two females using Canu [62] with the parameters “corOutCoverage=1000, minReadLength=1000, and correctedErrorRate=0.085”. (2) Sequencing errors were repaired using Arrow (https://github.com/PacificBiosciences/GenomicConsensus/) with the default parameters (only the females) and the Illumina paired-end reads obtained with Pilon [63] to increase the accuracy of the reference assembly. (3) The improved contigs were further rebuilt into two subassemblies (ref and alt) with HaploMerger2 [64]. (4) Based on the reference subassembly, clean Hi-C reads were analysed using Juicer v1.6.2 [65], and 3D-DNA [66] was then used to scaffold the contigs into pseudomolecules.

### Validation of the genome assembly

The genome completeness from the contig to chromosome-level assemblies was assessed using BUSCO v3.0217 [67]. The completed assembly was compared to the Plantae BUSCO “Embryophyta odb9” database, which contains 1,440 protein sequences and orthogroup annotations for key clades, using the default parameters. This result was then compared with that obtained for *M. alba* genomes (Table S2).

Furthermore, HISAT2 [68] was used to align the RNA-seq data to the two wild mulberry genomes, and the results showed 98.07% and 98.15% single-base mapping accuracy for *M. notabilis* and *M. yunnanensis*, respectively. We used BWA v0.7.8 [69] to map Illumina reads from short-insert-size libraries back to genome assemblies. Our findings indicated that 97.90% of the reads from *M. notabilis* and 99.50% of the reads from *M. yunnanensis* were mapped to the assemblies, implying that the assemblies were highly complete (Table S2).

The Hi-C heat map revealed a well-organized interaction contact pattern along the diagonals within each pseudochromosome (Figure S1A–B). The LTR-RT assembly index [14], a metric used to evaluate the completeness of a genome assembly based on the quality of the assembly of repeat sequences, was also used for all the abovementioned genomes using the LTR_retriever pipeline [70].

### RNA-Seq and transcriptome assembly

RNA-Seq data were generated from six tissues (root, bark, stem, male flower, female flower and leaf), and total RNA was extracted using RNAiso Plus (Takara, Dalian, China) according to the manufacturer’s protocols.

The Illumina HiSeq XTen platform was used to create and sequence 15 paired-end libraries containing sequences with a 150-bp read length, and Trimmomatic v0.36 [71] software was used to trim the adapter sequences of the RNA-Seq reads. Genome-guided transcriptome assembly was performed with HISAT2 and StringTie v1.3.475 [72]. HISAT2-build was used to construct the genome index, and HISAT2 was used to map the clean transcriptome reads to the *M. notabilis genome*. The findings were integrated with StringTie in merge mode after the transcripts for each sample were assembled. HISAT2 and the StringTie pipeline were used to calculate reads per kilobase per million (RPKM) values. The R package ‘edgeR’ [73] was used to investigate differentially expressed genes.

### Repeat annotation and gene prediction

A combination of homology searching and *ab initio* prediction was used to identify the repetitive sequences in the mulberry genome. We searched against Repbase with RepeatMasker [74] and RepeatProteinMask for homology-based prediction. We employed Tandem Repeats Finder [75], LTR FINDER [76] and RepeatScout [77]with default parameters for *ab initio* predictions.

The predictions of protein-coding genes were performed using previously reported methods with minor revisions [5]. Briefly, using Augustus [78], GlimmerHMM [79], and SNAP [80], we performed an *ab initio* coding region prediction in the repeat-masked genome. PASA-H-set gene models trained Augustus, SNAP, and GlimmerHMM, which were then used to predict three masked mulberry genomes. The annotated proteins from *M. notabilis*, *M. alba*, *C. sativa*, *Fragaria vesca*, *Malus domestica*, *Prunus persica*, *A. thaliana*, and *O. sativa* were used to obtain protein evidence. Next, we used Trinity to assemble the transcriptome based on RNA-Seq data and then PASA [81] to align the assembled sequence to the genome for gene predictions. Additionally, TransDecoder (https://github.com/TransDecoder) was also used to identify putative coding regions in transcript sequences. EVidenceModeler [82] was then used to combine the aforementioned findings to forecast the complete set of nonredundant genes. All predicted proteins were annotated using InterProScan v5.35-74.0 [83] and by running a BLASTP [84] search against the KEGG [85], Swiss-Prot and TrEMBL [86] databases with an E-value threshold of 1e^-5^.

### Identification of LTR-RTs

A comparative analysis of LTR-RTs was performed using the genome sequences of two *M. notabilis* (Male and Female), *M. yunnanensis* and *M. alba*. LTR-FINDER [76] (parameters -w 2 -d 0 -l 100) was used to detect LTR-RTs.

### Estimation of insertion time of the LTR-RTs

All LTRs sequences with complete 5′-LTR and 3′-LTR were used. MUSCLE [87] (with default parameters) was used to align the 5′-LTR flanking and 3′-LTR flanking sequences, and the distance between the alignment sequences was calculated using disMat (http://emboss.sourceforge.net/, with parameters -nucmethod 2). *T* = *K*/*2r* (divergence between LTRs/substitution per site per year) was used to calculate the insertion time. The mutation rate (per base per year) used was 1.8 × 10^−8^.

### Chromosome evolution

We used the genomes of *M. notabilis*, *M. alba*, *F. hispida*, *F. microcarpa*, and *C. sativa* (*C. sativa* as the outgroup) to reconstruct the ancestral chromosome karyotype of the Moraceae family based on a previously published technique with minor revisions [88]. Briefly, using *M. alba* as the reference genome, we performed pairwise alignments with other species as targets using LAST v1.1 with the default parameters. Subsequently, axtChain, chainMergeSort, chainPreNet, and ChainNet were used to generate “chain” and “net” files as inputs for DESCHRAMBLER (https://github.com/jkimlab/DESCHRAMBLER). We then identified 455 conserved segments using DESCHRAMBLER at a 1,200-kb resolution and reconstructed 21 predicted ancestral chromosomes with a total length of ∼ 305 Mb (Table S5).

### Comparative genomics and detection of EFRs in mulberry genomes

The MCscan toolkit (https://github.com/tanghaibao/jcvi/wiki/MCscan) was used to identify homologous gene pairs between the genomes of *M. notabilis*, *M. yunnanensis*, and *M. alba*.

We estimated large-scale homologous synteny blocks (HSBs) in the pairwise whole-genome alignment utilizing the chromosomal sequences of the *M. notabilis* and *M. alba* genomes to detect probable EFRs. Raw local synteny blocks between the two genomes were detected using the MCscan toolkit. An EFR was defined as the interval between two large-scale HSBs demarcated by the end-sequence coordinates of large-scale HSBs on each side. The relative gene density, SD content, and repetitive content within the EFRs of each chromosome were compared to the complete chromosome using an in-house script.

### Analysis of gene families and phylogenetic evolution

Orthologous gene families in *M. notabilis*, *M. yunnanensis*, *M. alba* and five other species (*V. vinifera*, *P. persica*, *C. sativa*, *F. hispida* and *F. microcarpa*) were identified based on annotated genes using OrthoFinder v2.2.7 [15]. The expansion and contraction of the *M. notabilis* and *M. yunnanensis* gene families were examined using CAFÉ [89]. Single-copy orthologous genes were subsequently extracted, aligned with MUSCLE v8.2.10 [87], and analysed phylogenetically using RAxML v8.2.10 [90] with the GTRGAMMA model. The species divergence time was estimated using MCMCTree in PAML v4.8 [91], and calibration times were determined using the TimeTree database (http://www.timetree.org/).

Four-fold degenerate sites (4DTv) from the whole-genome SNP collection were extracted and concatenated into a “supergene” format for each species to construct a *Morus* genus tree. The seven aligned 4DTv supergenes were used to construct a phylogenetic tree using the IQ-TREE [16] program.

### Detection of SNPs, small indels, SVs and SDs

SNPs and small indels (length ≤ 500 bp) were compared between the two wild genomes assembled in this work and the *M. alba* genome using MUMmer v3.2394 [92]. First, we used the number from MUMmer with the parameter “-mum -g 1000 -c 90 -l 40” to generate the alignment. The files were then filtered using the delta-filter comparison with the query “-r -q” to create a one-to-one map. Show-snp (a module of MUMmer) with the parameter “-ClrTH” was used to call SNPs and small indels from the one-to-one alignment module. Furthermore, the web-based SV analysis tool Assemblytics [93] was used to analyse large SVs. In addition, we identified SDs in the mulberry genomes with reference to the *Ficus* genomes (https://github.com/tangerzhang/popCNV).

### Identification of SDRs in *M. notabilis*

A previously described technique based on k-mers [19, 20] was used to identify sex-determining genes in the *M. notabilis* genome, with a slight modification. Using Jellyfish [58], the reads from male and female *M. notabilis* (four individual replicates per sex) were catalogued into 40-bp k-mers. Starting with an “AG” dinucleotide, their total cumulative counts in females and males were greater than ten, and the reads were defined as valid k-mers. In addition, k-mers with a count of zero in the female group were defined as MSKs, and those with a count of zero in the female group were defined as FSKs. We complemented this approach by mapping the sex-specific reads to the assembled male *M. notabilis* genome using BWA-MEM with the default parameters, and we prioritized genomic regions (10-kb windows) with a high depth of sex-specific reads. After deleting duplicates, SAMtools v1.9 [94] was used to calculate the mapping depth of the reference genome (by parsing specific 50-kb windows). Additionally, deduplicated BAM data were normalized using the deepTools bamCoverage tool [95].

### Quantifying the digital expression and analysing the potential function of candidate sex genes

Illumina sequencing experiments (Illumina NovaSeq 6000, USA) were performed at the levels of lncRNA, small RNA and DNA methylation to quantify the digital expression and to explore the gene regulation patterns of candidate sex genes. Libraries were constructed using a NEBNext^®^ Ultra™ RNA Library Prep Kit (#E7530L, NEB, USA) (mRNA), TruSeq Stranded Total RNA Library Prep Kit (lncRNA), TruSeq Small RNA Library Preparation Kit (small RNA), and TruSeq Methyl Capture EPIC Library Prep Kit (DNA methylation), respectively, according to the manufacturers’ instructions.

In the analysis of RNASeq, lncRNASeq, and sRNASeq data, rRNA/tRNA contaminants were removed by mapping the reads to rRNA/tRNA sequences from public databases. Transcripts of lncRNAs were assembled using Trinity V2.6.6 with the parameter settings for strand-specific reads (‘--SS_lib_type RF’). The obtained sequences were used to search the NCBI NR database. Small RNA reads and bisulfite sequencing (BS-Seq) reads were mapped to the reference genome using Bowtie2 v2.3.4.1 and Bismark v0.16.3 [96], respectively. The alignment results were visualized using Integrative Genomics Viewer (IGV) v 2.4.14 [97].

### Read alignment and variant calling

The filtered reads from all individuals were aligned to the *M. alba* genome using BWA-MEM. The read pairs were filtered using Picard tools v2.18 (https://broadinstitute.github.io/picard/). SNPs were identified in all the samples using the HaplotypeCaller module of GATK v3.868 [98] in genomic variation call format (gVCF) mode. Briefly, HaplotypeCaller was used to identify the gVCF for each sample. Subsequently, all gVCF files were merged to create a raw population genotype file with the SNPs. SNPs were preliminarily filtered using the GATK VariantFiltration function with the parameter “QD < 2.0 || FS > 60.0 || MQ < 40.0 || MQRankSum < -12.5 || ReadPosRankSum < -8.0 || SOR > 3.0 and mean variant sequencing depth (including all the individuals) < 1/3× and > 3×”. In addition, SNPs were filtered according to the following criteria: (1) MAF ≥ 5%, (2) maximum missing rate < 0.1, and (3) restriction to two alleles.

### Population structure and phylogenetic analysis

The NJ tree, PCA and ADMIXTURE methods were used to explore the genetic relationships among wild and domesticated mulberry populations. An NJ tree was constructed for the whole-genome SNP set using MEGA v6.0 [99] based on a pairwise genetic distance matrix, which was calculated using PLINK v1.9 [100] with the option “--distance-matrix”. PCA was performed using SmartPCA in EIGENSOFT v6.1 [101]. The significance of eigenvectors was assessed using the Tracy–Widom test. Population genetic structure was inferred using ADMIXTURE v1.3.0 [102] considering k = 2 to k = 10 (Figure S11), and the analysis was repeated 20 times for each k value. For the PCA and ADMIXTURE analysis, we used a nonredundant SNP data set obtained after removing rare alleles with the option “--indep-pairwise 50 5 0.4” in PLINK and further excluding SNPs with intrachromosomal LD r^2^ values < 0.4 to remove the bias caused by LD.

The value of π was calculated using VCFtools v0.1.15 [103] based on the high-confidence filtered SNPs. The π value for each SNP was calculated, and the nucleotide diversity level was measured using “--window-pi 50000 --window-pi-step 20000” for each subpopulation. LD decay was calculated for all pairs of SNPs within 500 kb using PopLDdecay v3.27 [104] with the default parameters.

TreeMix (version 1.13) was used to infer models of population split and migration between groups. TreeMix was run using the allele frequencies calculated from the LD-pruned SNP set with the parameters “-bootstrap 5000 -global” and “migration event -m” (from 0–4).

### Identification of selective sweeps

We performed the analysis described below to detect selective sweeps during mulberry domestication. (1) The high divergence in genetic diversity (π ln ratio) and high fixation index *F*_ST_ were analysed by parsing specific 50-kb windows between the domesticated group (JP, GD, HU, and NH) and WD. (2) Putative selective sweeps were defined as windows with outlier signals (top 1%) overlapping for the two statistics (*F*_ST_ > 0.564, π ln-ratio WD/JP-GD-HU-NH > 2.041). (3) Tajima’s D and comparison haplotypes were applied to confirm the top signals.

### Functional enrichment analyses

We characterized the most relevant functions of the protein-coding genes with chromosomal break regions and selective sweeps by searching for overrepresented KEGG pathways and GO terms. Target protein sequences were used to conduct functional enrichment tests of the target genes using KOBAS 3.0 (http://kobas.cbi.pku.edu.cn/kobas3/annotate/). The *P* value was calculated using a hypergeometric distribution, and *P* values < 0.05 were considered to indicate significantly enriched pathways.

### RNA extraction and qRT–PCR analysis

Total RNA was extracted as described above, and cDNAs were generated with a PrimeScript RT Reagent Kit with gDNA Eraser (Takara). We performed qRT–PCR with TB Green Premix Ex Taq II (Takara). Relative expression was calculated using the 2^−ΔΔCt^ method [105]. The primers used for the gene expression analysis are listed in Table S9. The values are presented as the means ± SDs from three biological replicates (\*\**P* < 0.01, \*\*\**P* < 0.001, and \*\*\*\**P* < 0.0001), as determined using one-way analysis of variance (ANOVA).

### PCR validation of candidate sex-determining genes

Degenerate primers were designed for four candidate genes to verify their male specificity (Table S10). The PCR mix was composed of 10 μL of 2× Ex Taq MasterMix (CWBIO), 1 μL of each primer, and 1 μL of genomic DNA at a concentration of ∼100 ng/μL, and ddH_2_O was added to obtain a total reaction volume of 20 μL. The thermocycling conditions were as follows: an initial cycle of 5 min at 94°C, followed by 35 cycles of 30 s at 94°C, 30 s at 57°C, 30 s at 72°C, and 2 min of extension at 72°C. The PCR products were loaded on a 1.0% agarose gel and run at 150 V for 15 min. The samples were imaged under ultraviolet light.

## Data availability

The genome assemblies, gene annotation and Illumina re-sequencing short reads have been deposited to Genome Sequence Archive (GSA) [106] and Genome Warehouse (GWH) database in National Genomics Data Center, Beijing Institute of Genomics, Chinese Academy of Sciences/China National Center for Bioinformation (https://ngdc.cncb.ac.cn/gsa) under BioProject Accession number GSA: PRJCA008608.

## CRediT author statement

**Zhongqiang Xia:** Investigation, Data Curation, Data Analysis, Validation, Visualization, Writing - Original Draft, and Writing - Review & Editing. **Xuelei Dai:** Investigation and Data Analysis. **Wei Fan:** Investigation, Data Curation, and Writing - Review & Editing. **Changying Liu:** Investigation and Writing - Review & Editing. **Meirong Zhang:** Validation. **Peipei Bian:** Data Curation. **Yuping Zhou:** Validation. **Liang Li:** Validation and Visualization. **Baozhong Zhu:** Resources. **Shuman Liu:** Resources. **Zhengang Li:** Resources. **Xiling Wang:** Investigation and Resources. **Maode Yu:** Project administration. **Zhonghuai Xiang:** Project administration. **Yu Jiang:** Conceptualization, Writing - Original Draft, and Writing - Review & Editing, and Supervision. **Aichun Zhao:** Conceptualization, Resources, Writing - Original Draft, and Writing - Review & Editing, and Supervision. All authors have read and approved the final manuscript.

## Acknowledgements

This work was supported by the National Key R&D Program of China (Grant No. 2019YFD1000604), the China Agriculture Research System (Grant No. CARS-18-ZJ0201), the Forestry Promotion by Science and Technology Program of Chongqing, China (Grant No. Yulinkeyan2020–2) and Sichuan Science and Technology Program (Grant 22NSFSC3680).

## Supplementary material

**Figure S1.**
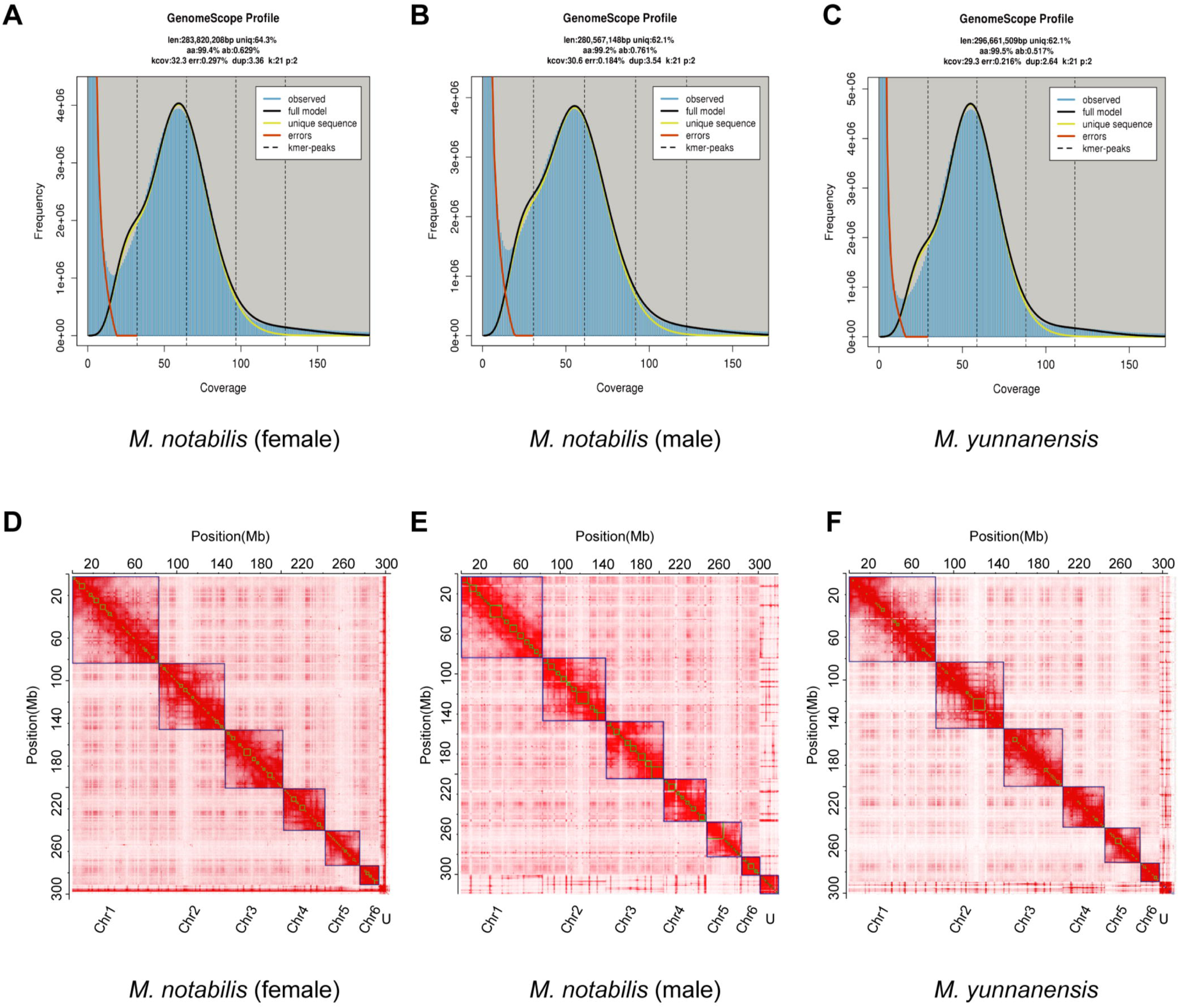
Genome size estimation and assessment of the three wild mulberry individuals with chromosome-level genome assemblies. **A**, **B** and **C**. Based on k-mer genome size estimates and visualized using online GenomeScope software. **D**, **E** and **F**. Chromatin interaction heatmaps of the genomes of dioecious *M. notabilis* and *M. yunnanensis*. U denotes an unanchored sequence.

**Figure S2.**
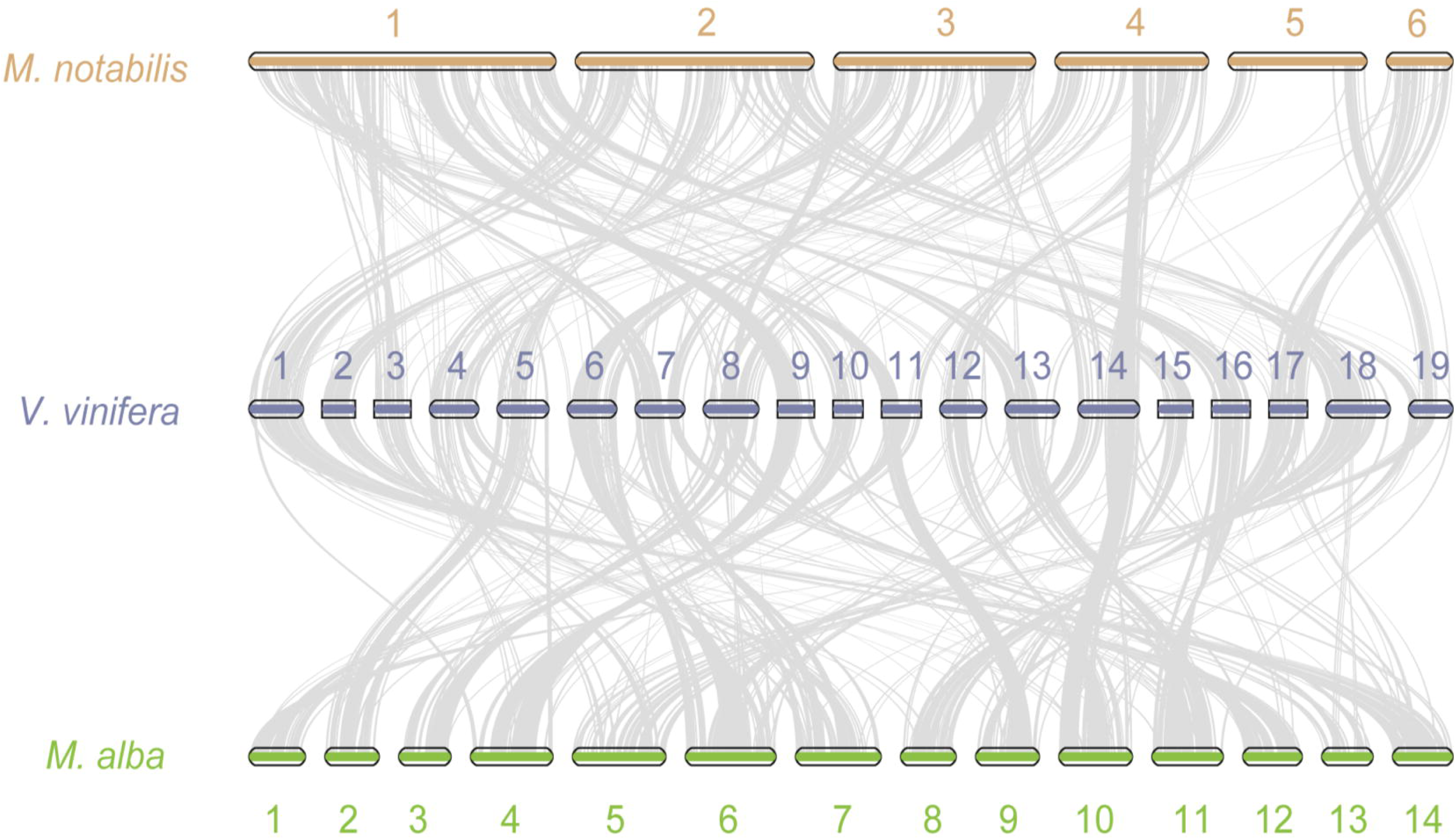
Synteny analysis of *M. notabilis* and *M. alba* with *Vitis vinifera*.

**Figure S3.**
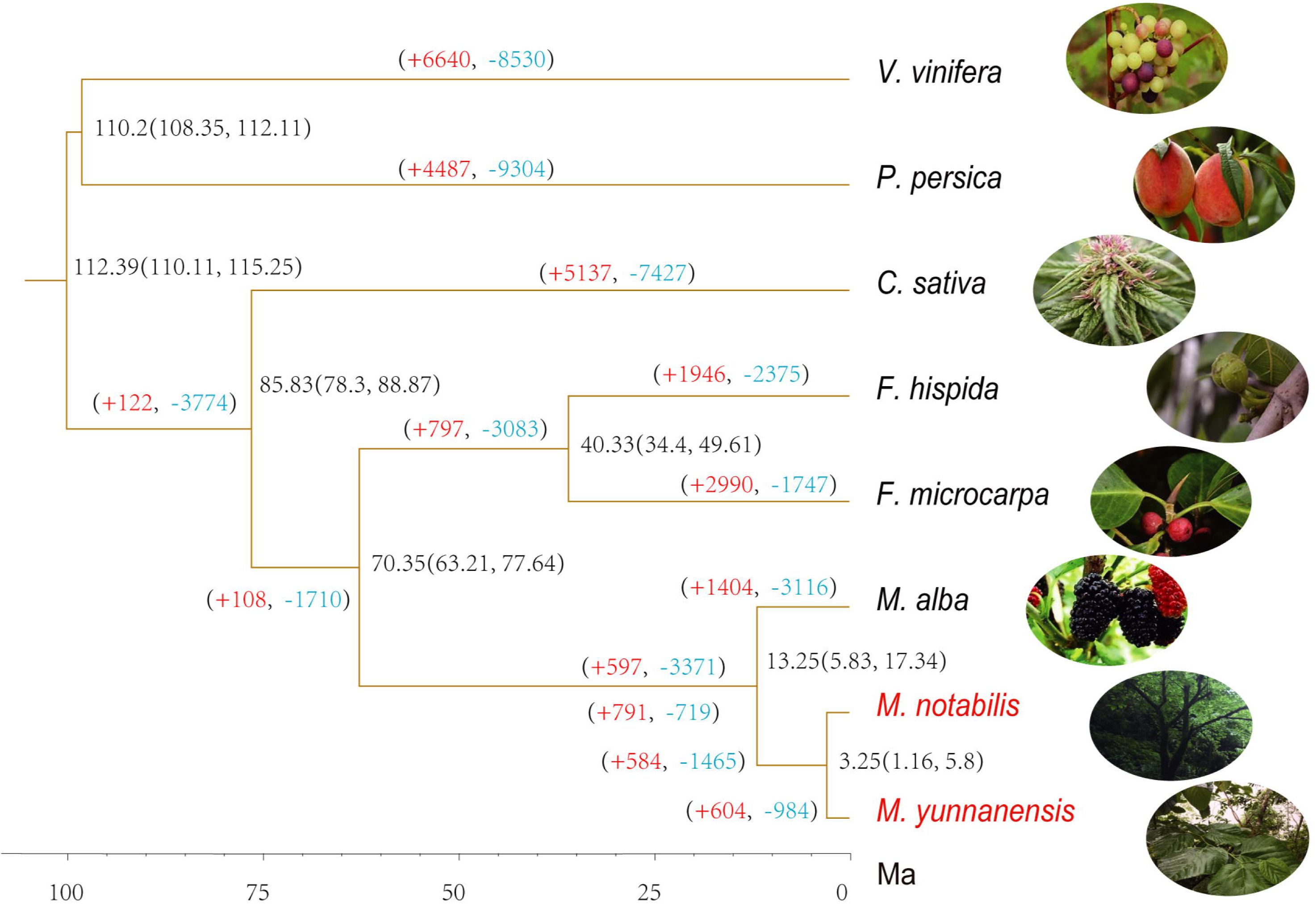
Phylogenetic relationships among *M. notabilis*, *M. yunnanensis*, *M. alba*, *V. vinifera*, *P. persica*, *C. sativa*, *F. hispida* and *F. microcarpa*. The divergence times among different plant species are labelled on the right. The numbers on each branch represent expansion (red) and contraction (blue) of gene families.

**Figure S4.**
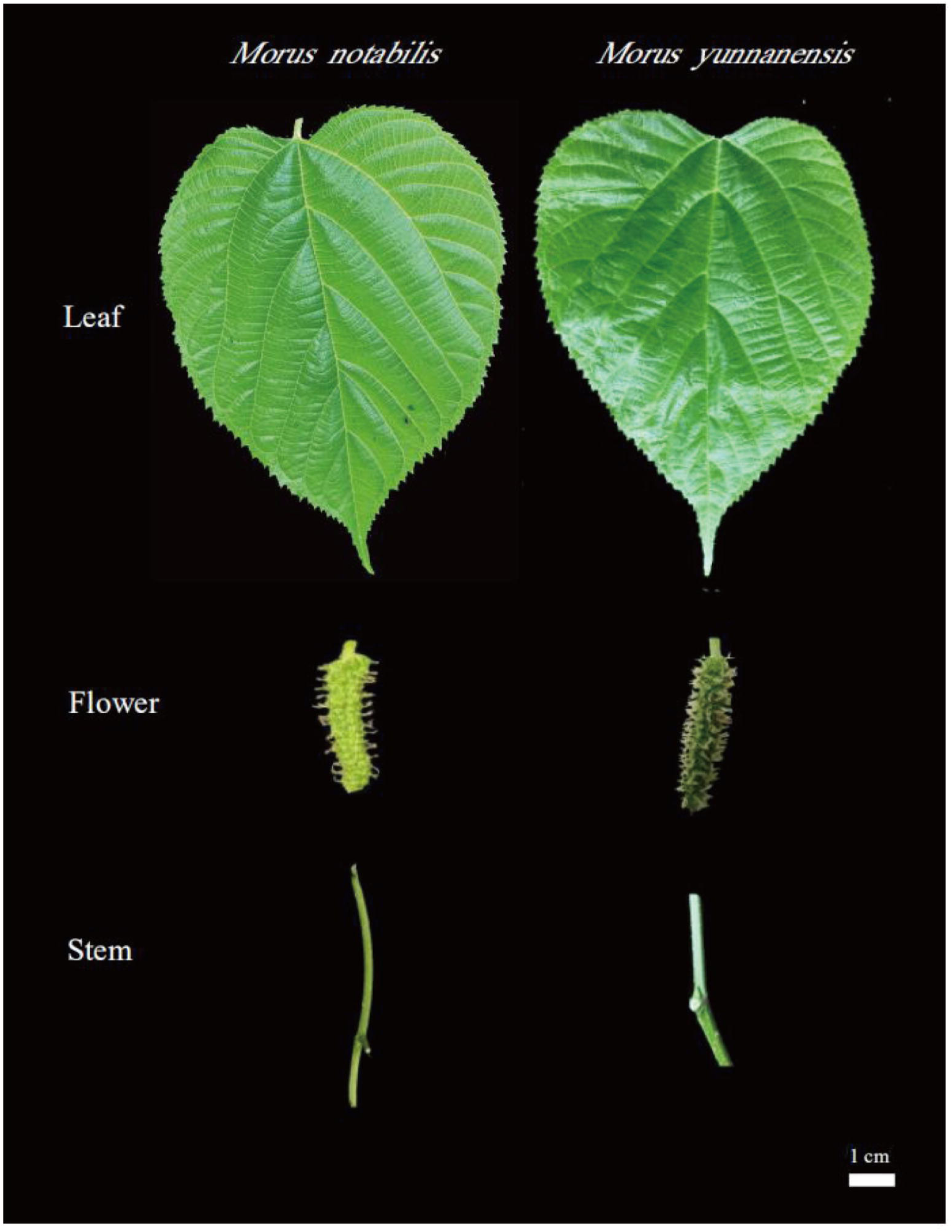
Leaf, flower, and stem organ morphology of the two de novo-assembled wild *Morus* species.

**Figure S5.**
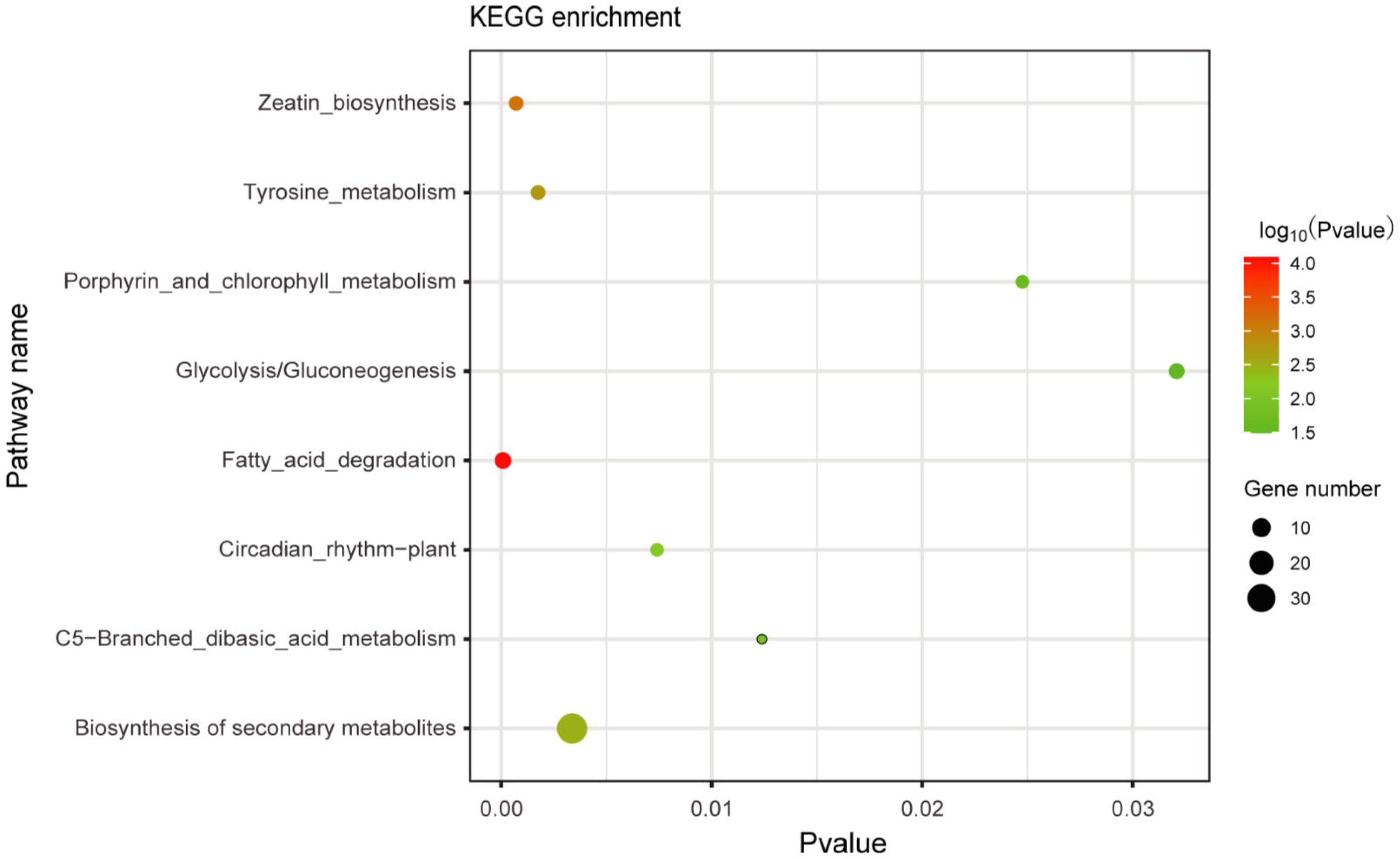
KEGG enrichment analysis of genes in segmental duplication (SD) regions in the *M. notabilis* genomes.

**Figure S6.**
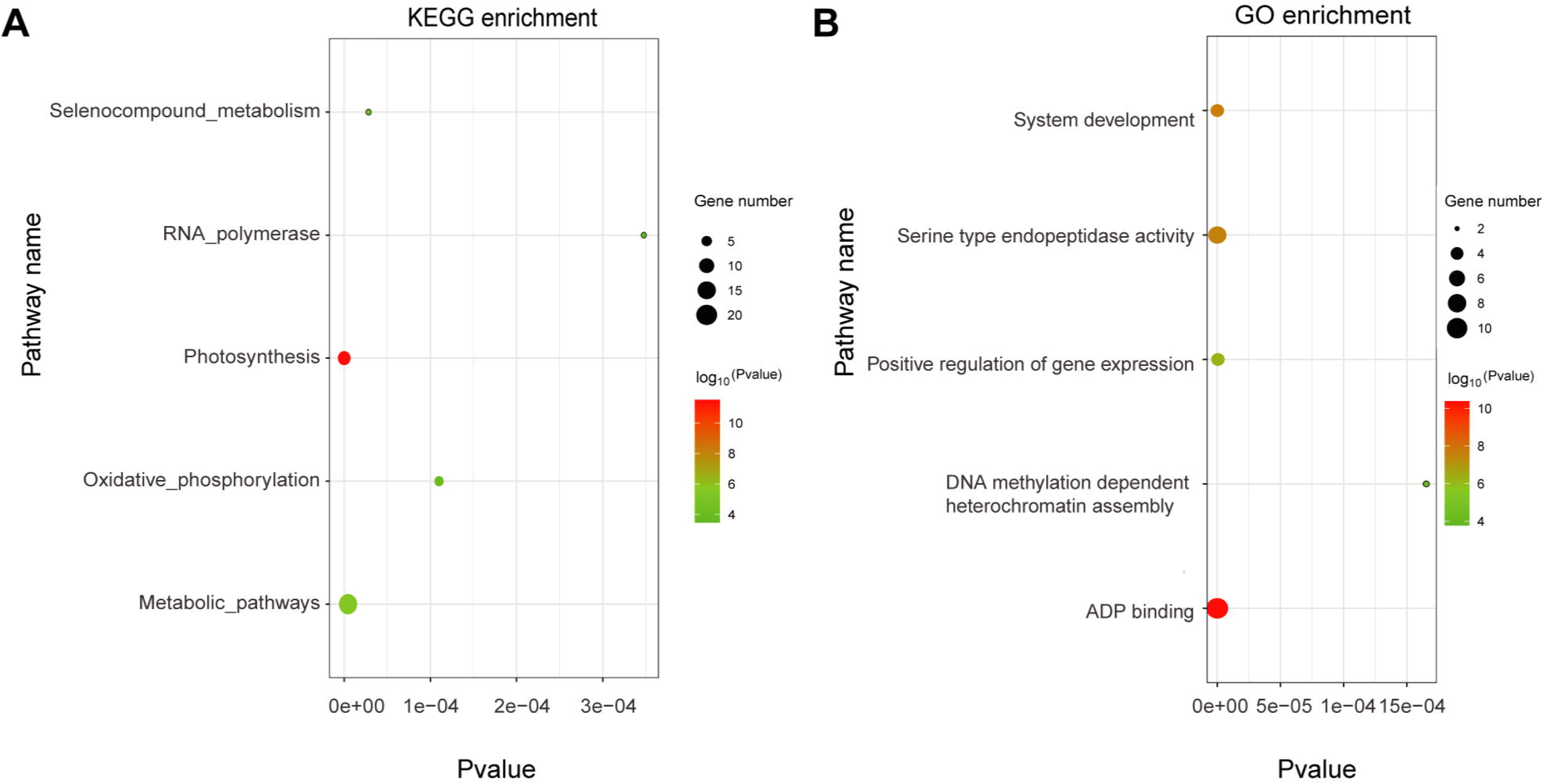
Functional enrichment analysis of genes in chromosome fusion regions in the *M. notabilis* genomes.

**Figure S7.**
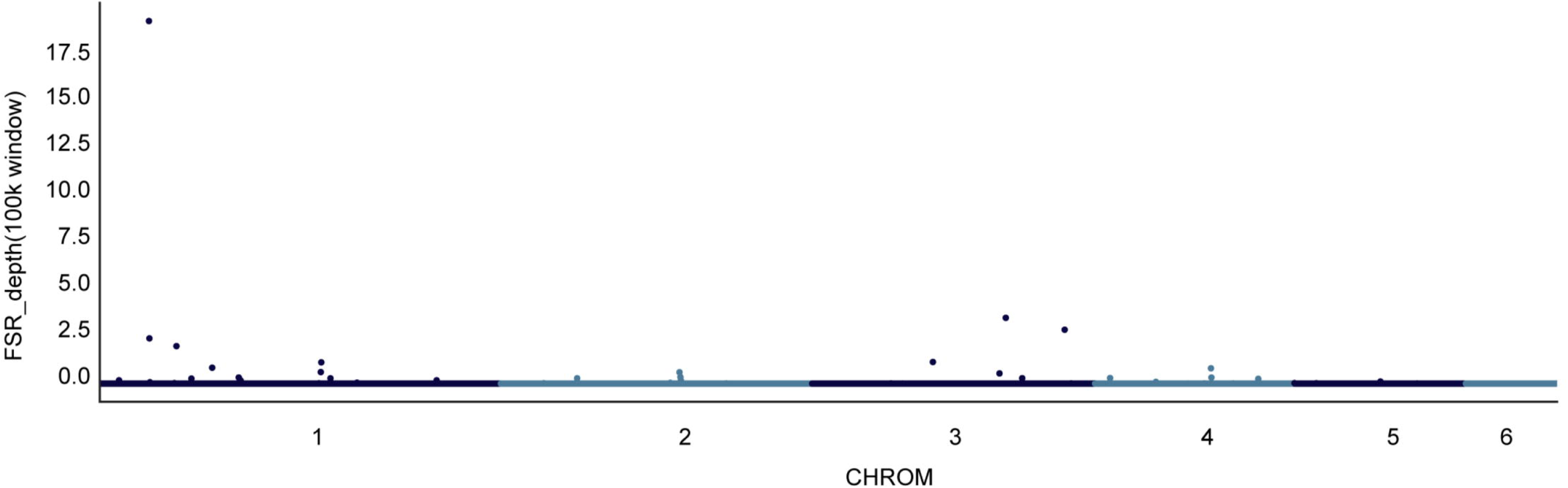
Manhattan plots of the mapping depths of female-specific reads (FSRs) in the male *M. notabilis* genome.

**Figure S8.**
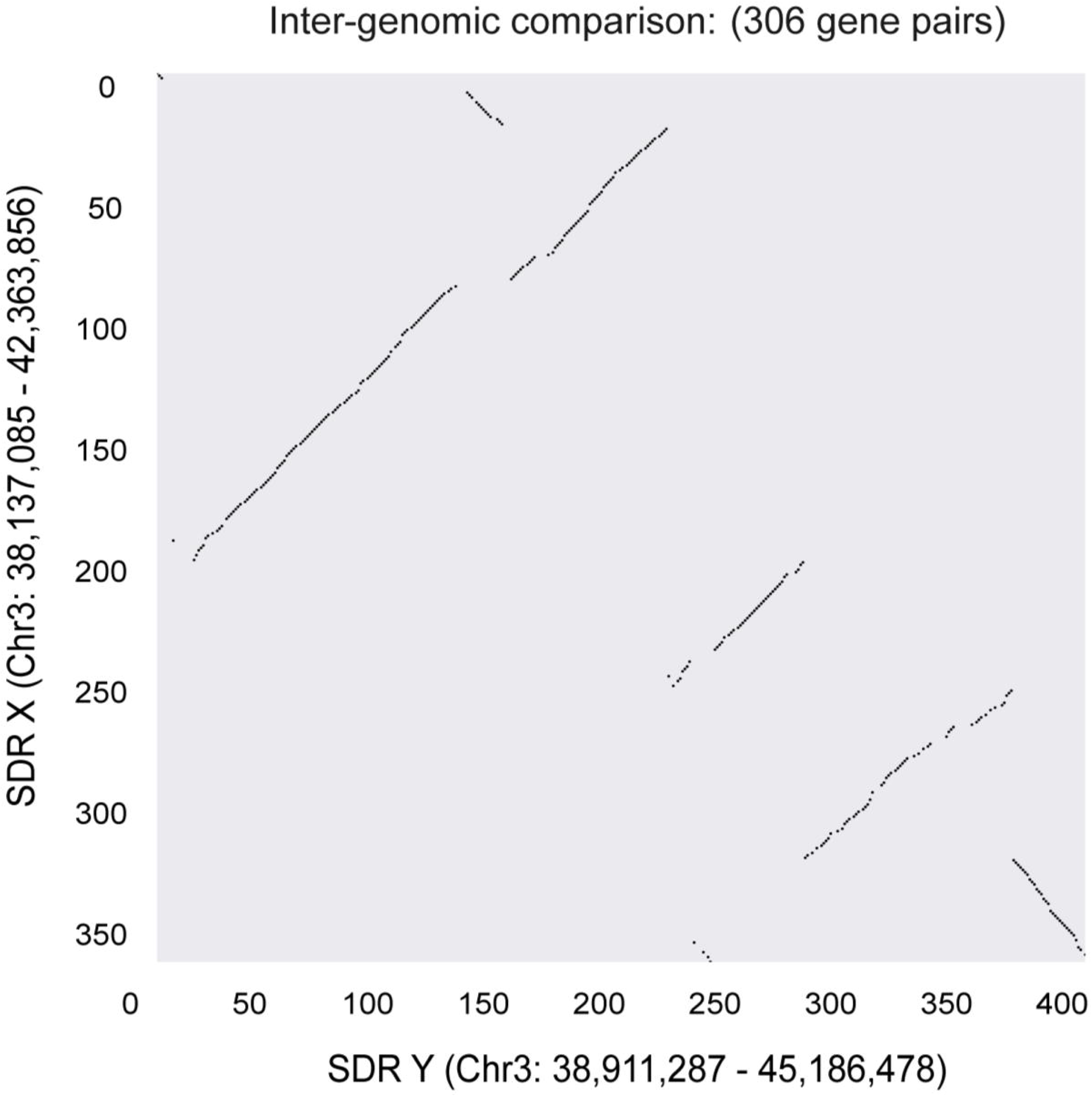
Genomic landscape of the SDR and its X counterpart. The black lines connecting gene pairs were identified using MCscan.

**Figure S9.**
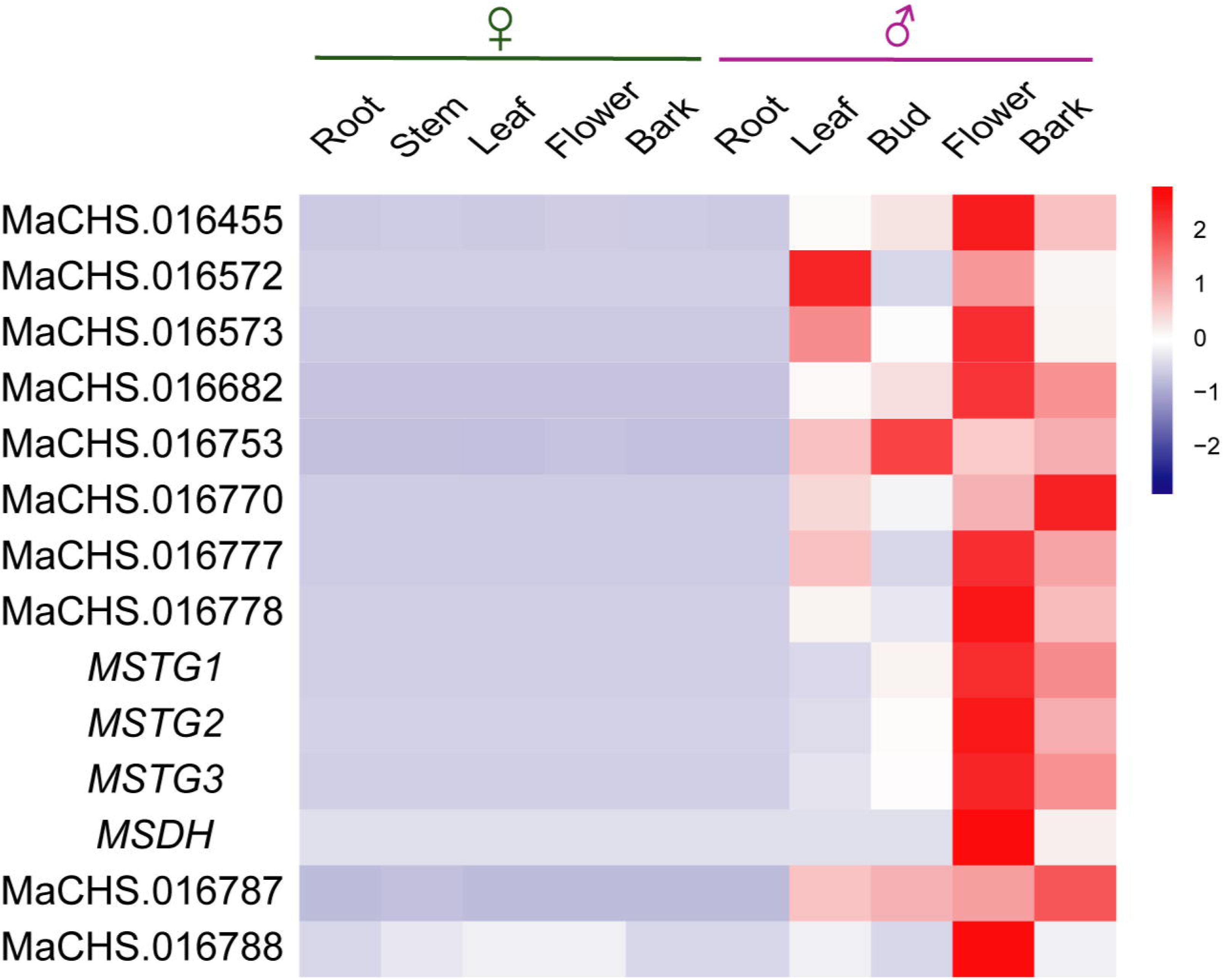
The expression of 14 candidate genes among unpaired genes that were significantly overexpressed in males was observed.

**Figure S10.**
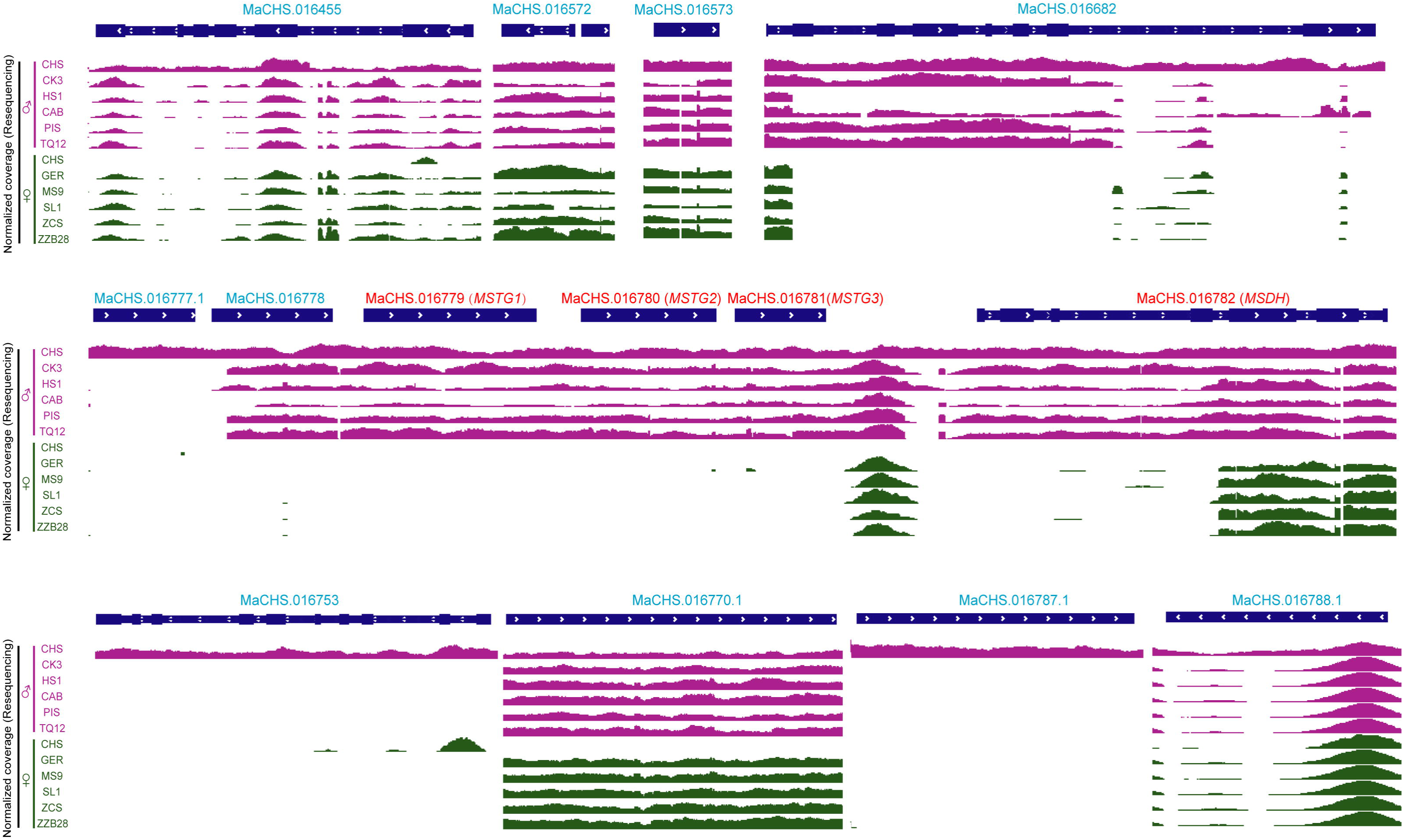
Alignments of male and female Illumina reads to the *M. notabilis* genome using IGV. The result revealed the presence/absence variants (PAV) three *MSTG*s and *MSDH* in male and female individuals, which confirmed them as candidate Y-specific genes.

**Figure S11.**
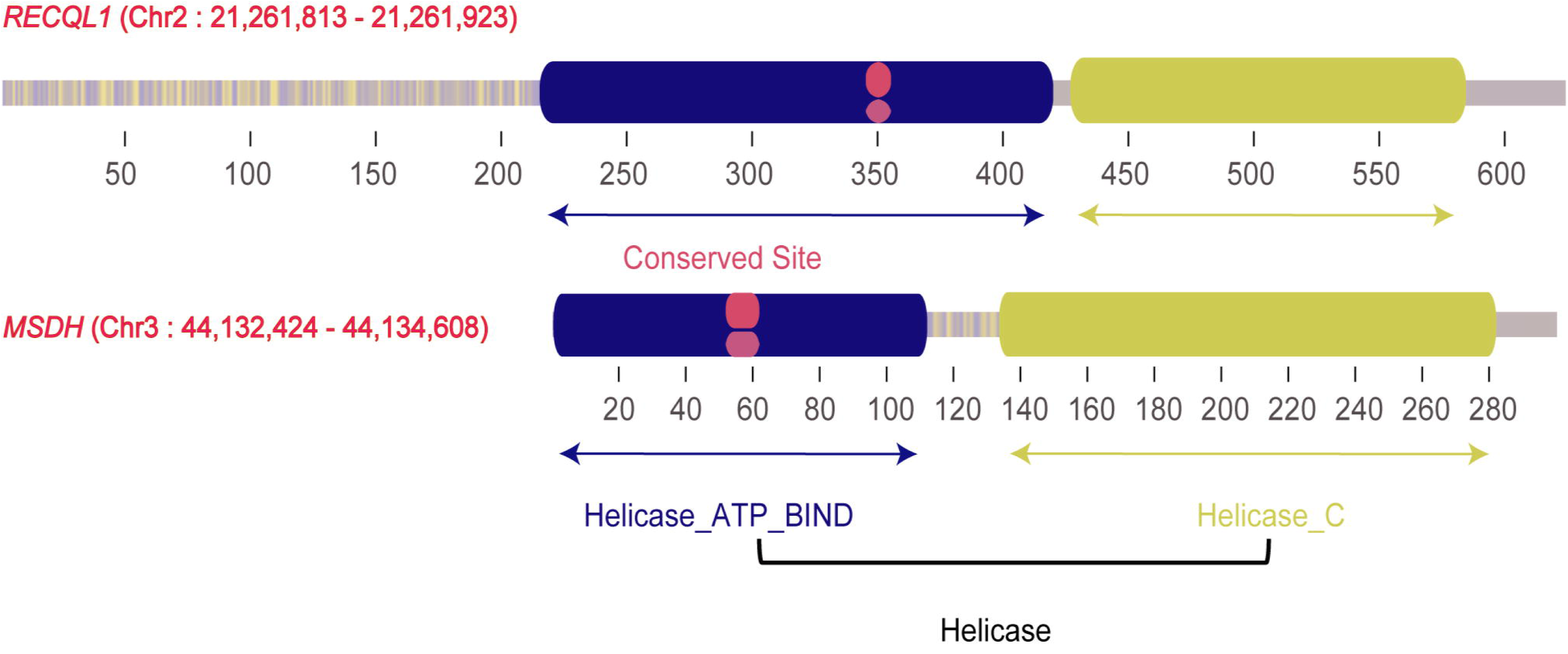
Homologous domains of *MSDH* and *RECQL1*.

**Figure S12.**
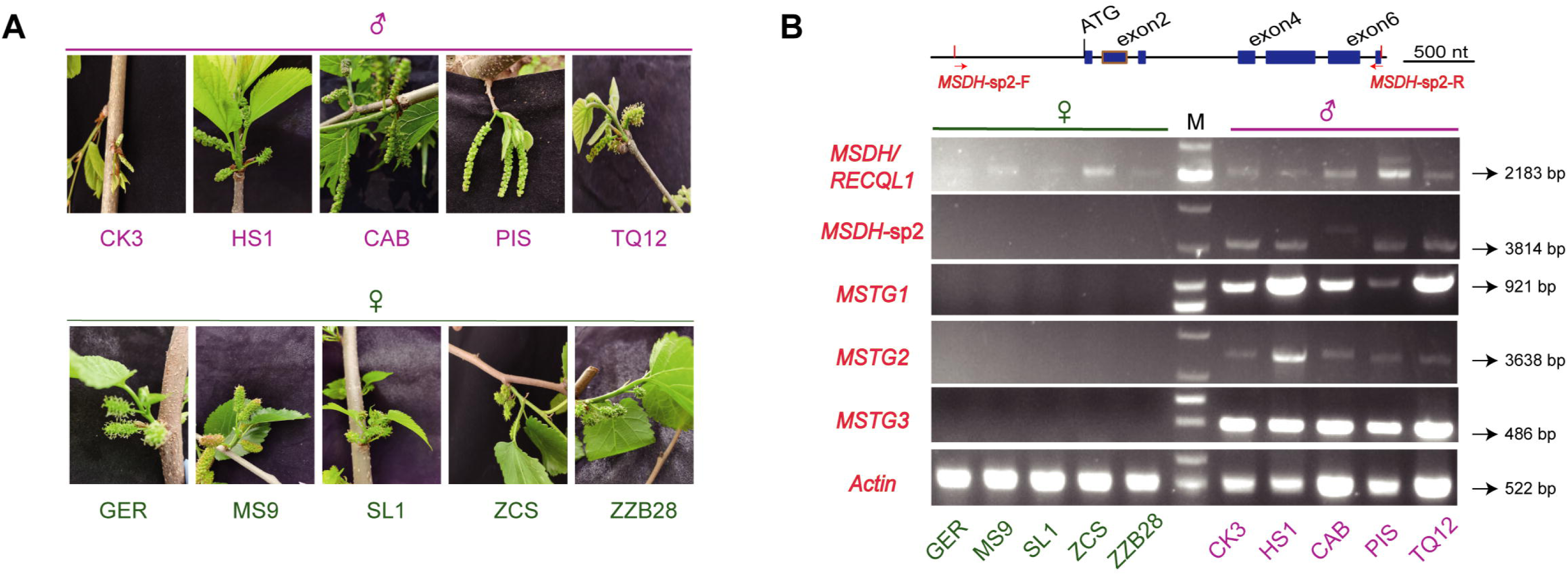
Validation of four candidate sex-determining genes in other dioecious *Morus* subgenera. **A**. Floral organ phenotypes of multiple diecious *Morus* species. GER: Georgia; MS9: *M mongolica* (Bur.) Schneid. Mengsang-9; SL1: *M. alba* L. Sri Lanka-1; ZCS: *M. alba* L. Zhichuisang; ZZB28: *M. alba* L. Zhenzhubai-2; CK3: *M. alba* L. Chenkou-3; HS1: *M. alba* L. Huosang-1; CAB: *M. australis Poir*. Cambodia; PIS: *M. alba* L. Pisang; TQ12: *M. alba* L. Tianquan-12. **B**. Agarose gel electrophoresis profile for four candidate sex-determining genes in different sexes corresponding to A. The location of the *MSDH*- specific2 (*MSDH*-sp2) primer is shown in top. Actin was used as the control. M, molecular marker. The location of the *MSDH*-sp2 primer is indicated by a red arrow. The primer sequences are listed in Table S10.

**Figure S13.**
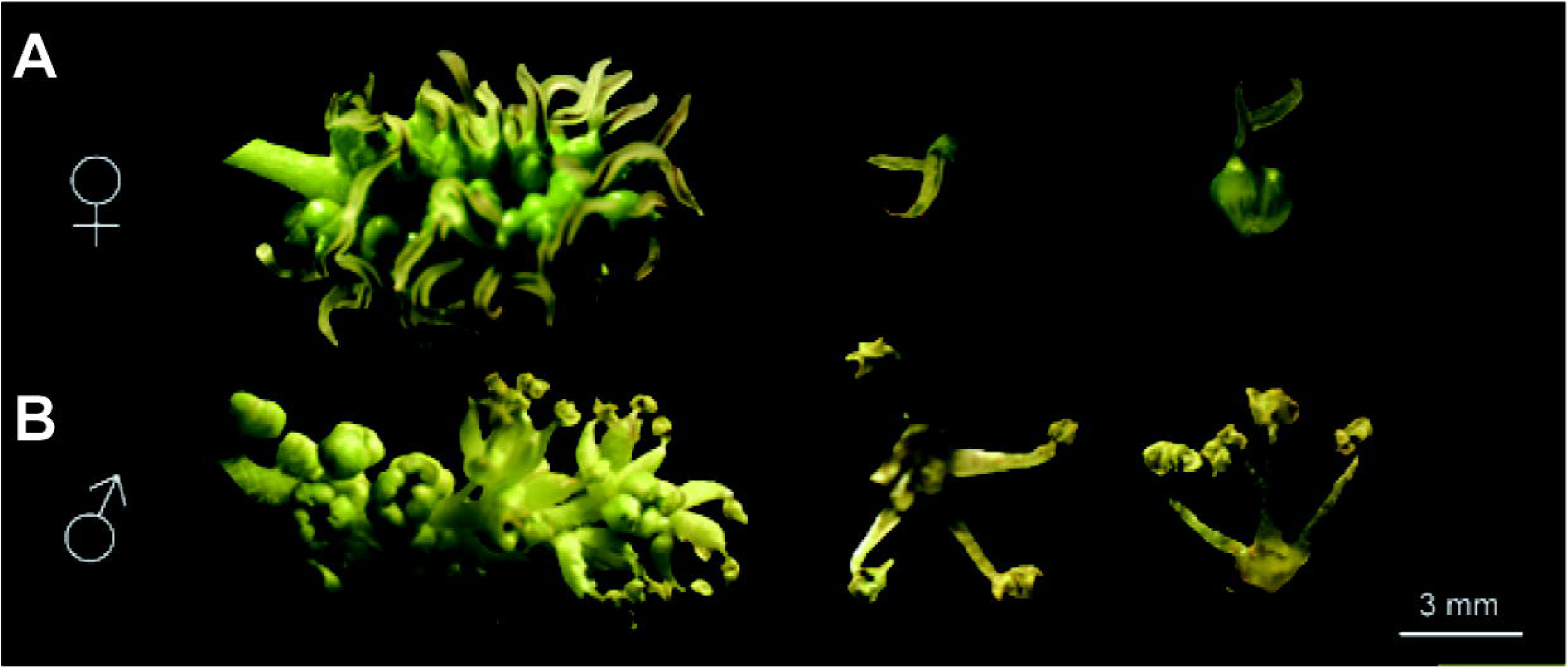
Morphology of the male and female flower buds defined in the text. The female flower morphological characteristics were photographed from female individuals (*M. alba* L. Zhenzhubai-2) and male individuals (*M. alba* L. Naxi). The photograph was taken by one of our co-authors, Dr. Wei Fan.

**Figure S14.**
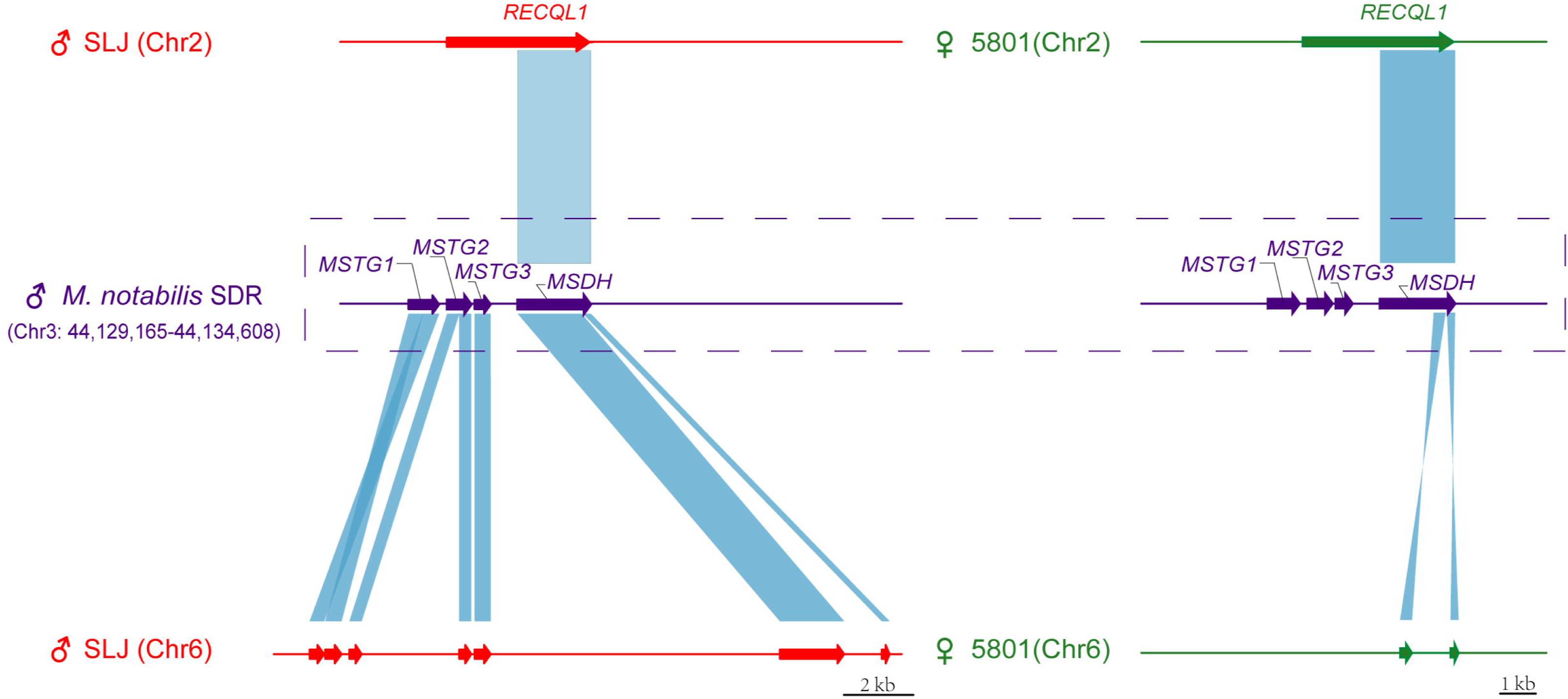
Schematic depicting the alignment of the candidate sex-determining locus to available genome assemblies of mulberry species. Red represents male, and green represents female. The top panel presents the structure of genes. SLJ (Male *M. wittiorum*. wild. SLJ, 2n=28, Unpublished data), 5801 (Female *M. alba*. cv. 5801,2n=28, Unpublished data).

**Figure S15.**
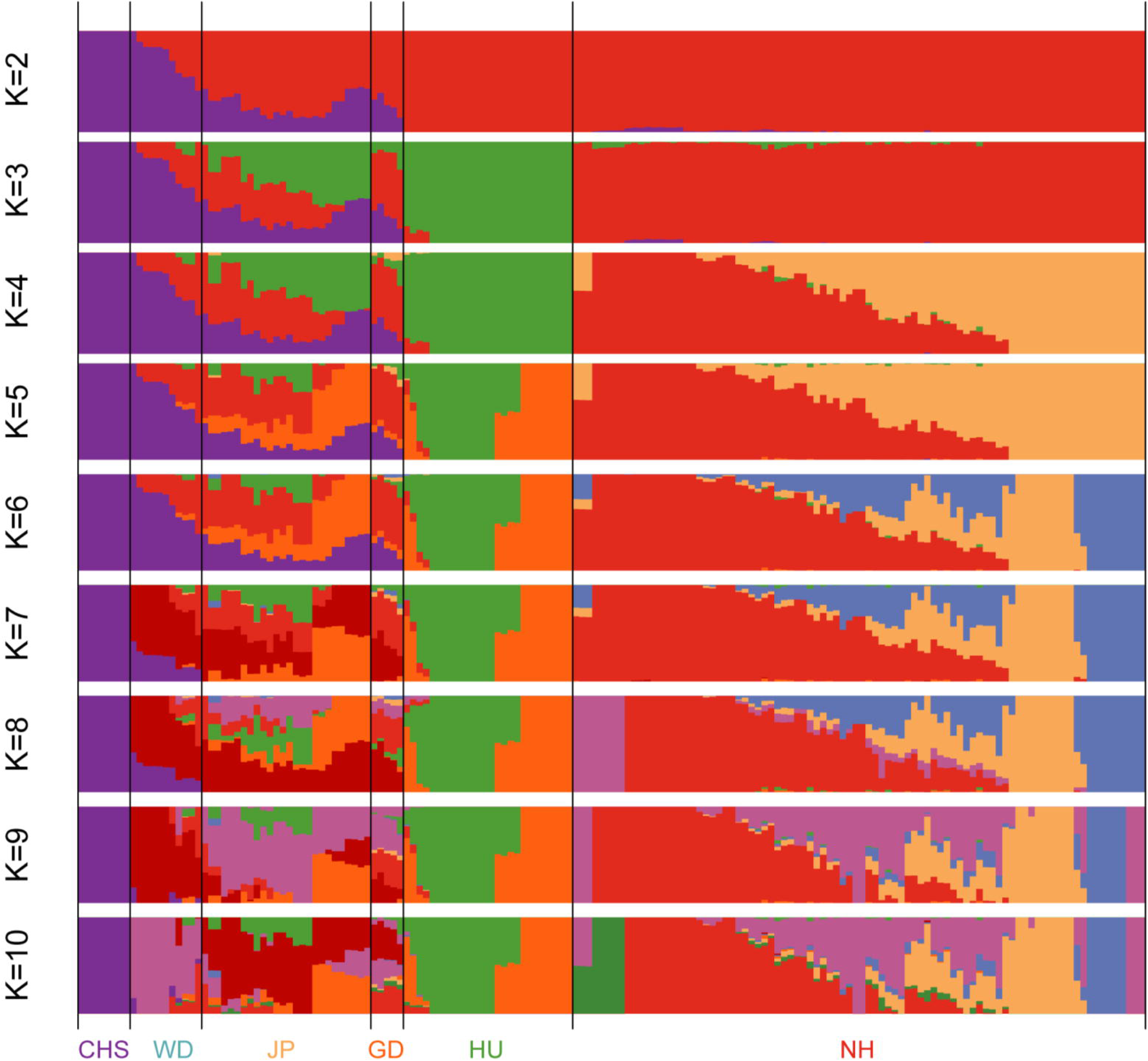
ADMIXTURE analysis from k = 2 to k = 10.

**Figure S16.**
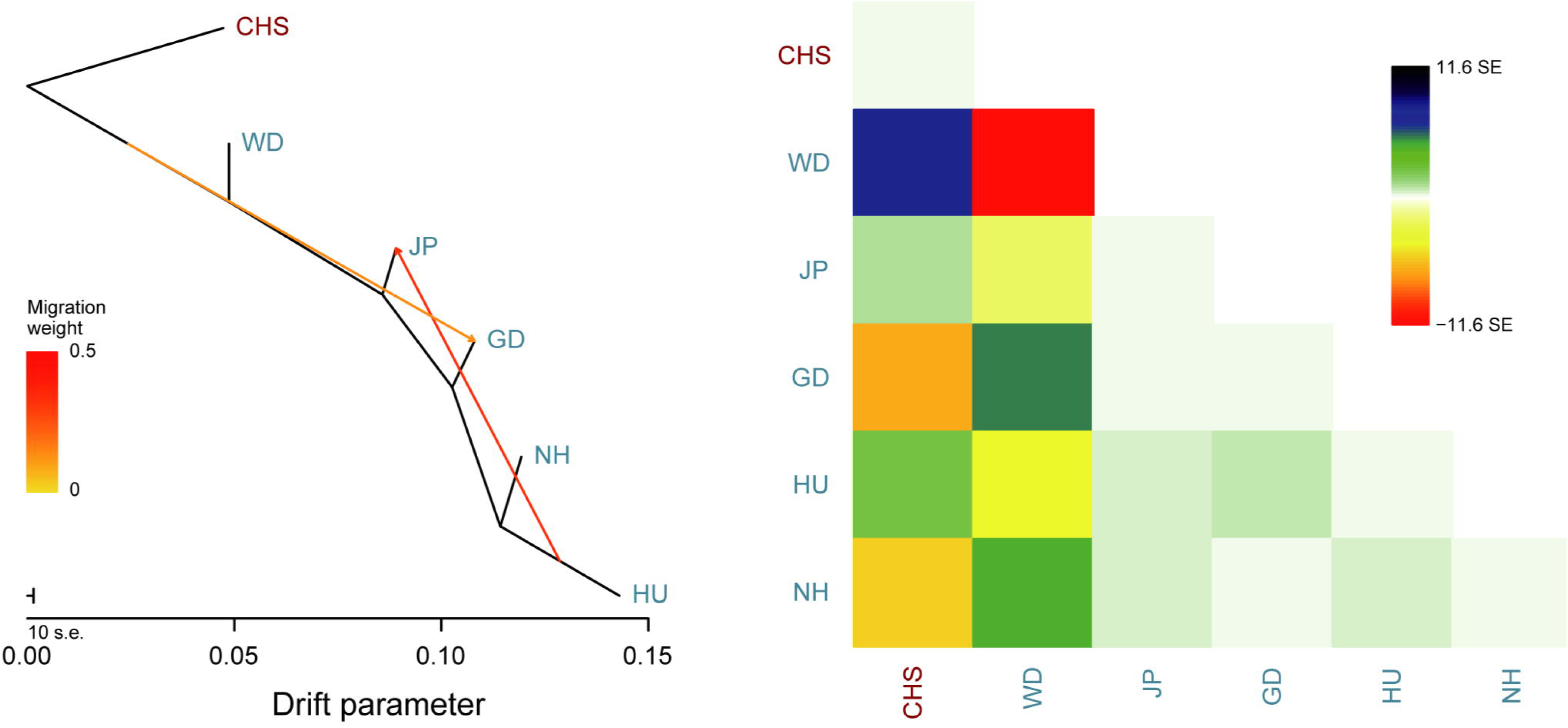
The TreeMix population splits between six “core” mulberry groups and the *M. notabilis* (CHS) group as the outgroup. **A**. Population relationships as inferred from the TreeMix analyses with 2 possible migration edges. **B**. Residuals of the covariance matrices.

**Figure S17.**
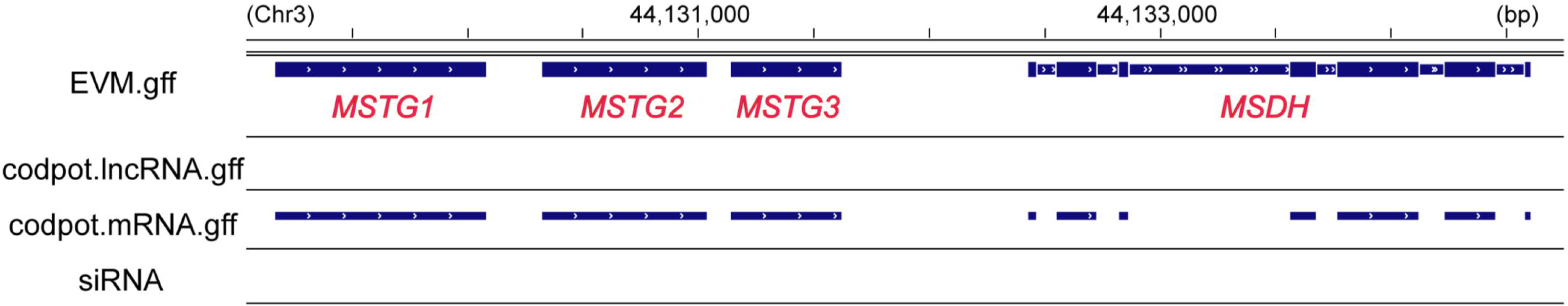
Validation of the siRNA abundance and protein-coding capacity of four candidate sex-determining genes in IGV. The top panel (EVM.gff) presents the structure of this gene. The panel ‘‘codpot.lncRNA.gff’’ indicates the assessment of the protein-coding potential of transcripts using lncRNA-seq reads, supporting alternative splicing sites for the gene structure. The panel ‘‘codpot.mRNA.gff’’ indicates supporting alternative splicing sites for the gene structure. The final panel ‘‘siRNA’’ includes small RNA-seq alignment of clean reads and indicates no small interfering RNAs of these genes.

**Figure S18.**
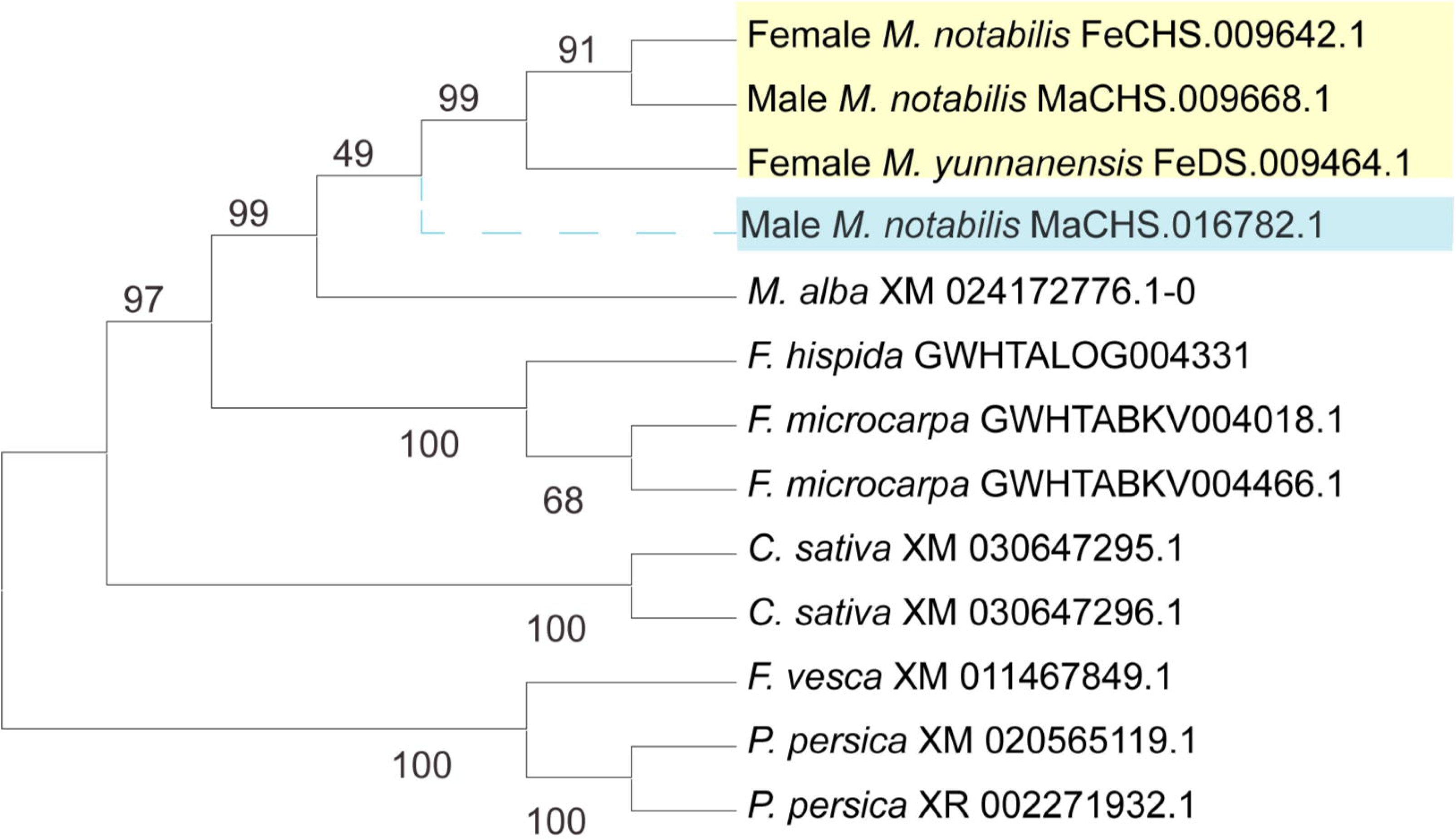
The duplication event of *MSDH* shown as a dashed line in the phylogenetic tree. Clades of *MSDH* and *RECQL1* are in blue and orange, respectively.

**Figure S19.**
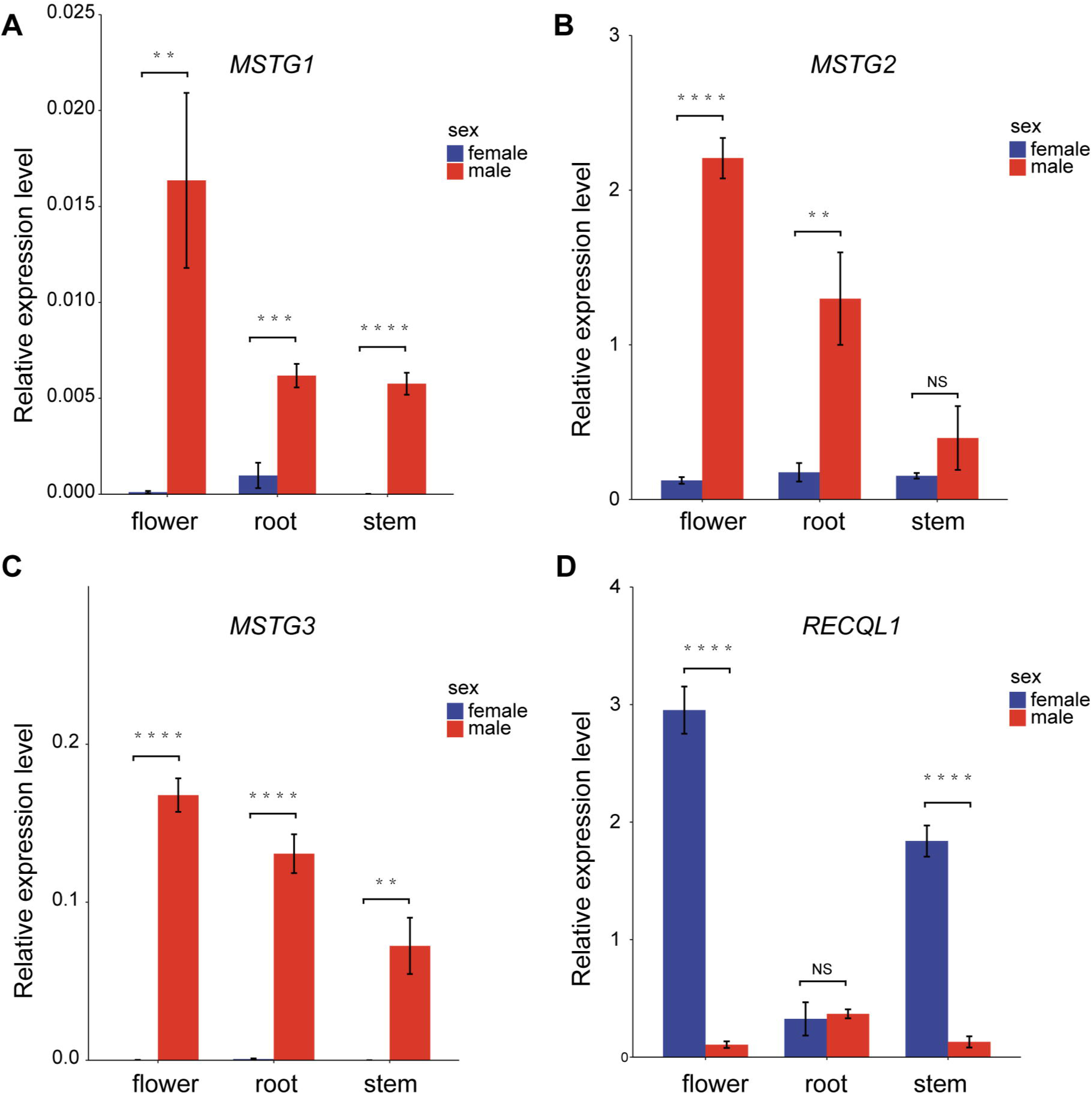
Validation of the digital expression of candidate sex determination genes in the flowers, roots and stems. The values shown are the means ± SDs from three biological replicates. Differences between groups were determined by one-way analysis of variance (ANOVA), **P < 0.01, ***P < 0.00, ****P < 0.0001. NS, not significant. The data are provided in Table S11.

**Figure S20.**
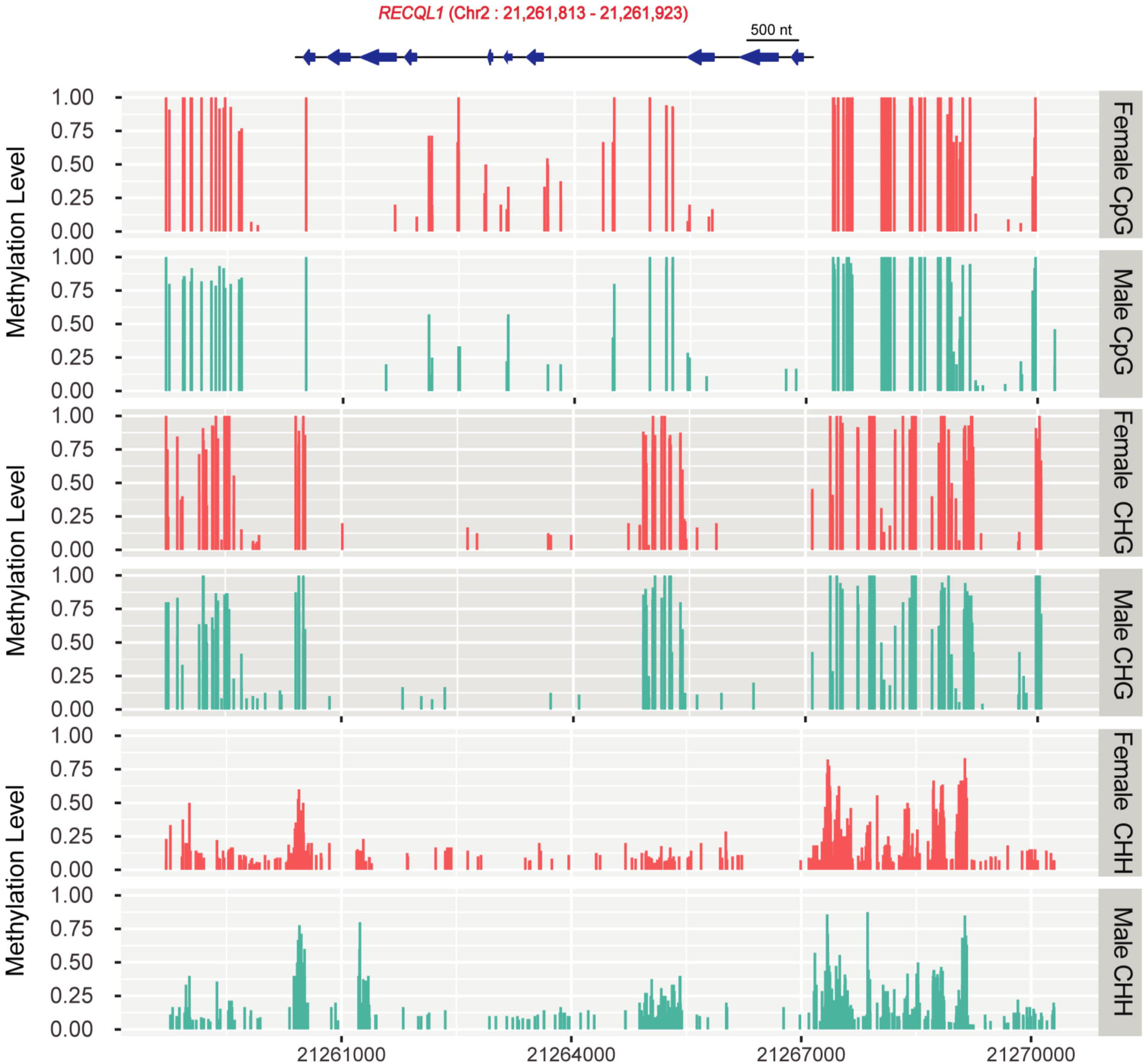
Differential methylation level of *RECQL1* in the two sexes.

## File Name: Supplemental Tables

### Description

**Table S1 Genome assembly statistics**

**Tables S2 Assessment of assemblies**

**Tables S3 Repetitive sequences**

**Table S4 Structural variations**

**Table S5 Statistics of conserved segments**

**Table S6 Characteristics of repeated sequences in chromosomal rearrangement regions**

**Table S7 Chromosome rearrangement region gene FPKM in different karyotypes**

**Table S8. Genes annotated on SDR-X and SDR-Y, including gene pairs identified using MCscan**

**Table S9 Statistics of transcriptome expression of the SDR gene in different tissues**

**Table S10 Gene amplification and quantitative specific primer sequences**

**Table S11 Tissue expression profiles of sex-candidate genes**

**Table S12 List and description of the mulberry accessions used in this study**

**Table S13 Population analysis of BAM statistical results**

**Table S14 SNP statistical results**

**Table S15 Cross-validation (CV) errors for ADMIXTURE ancestry models with k values ranging from 2 to 10**

**Table S16 Nucleotide diversity of different groups**

**Table S17 GO analysis of selected genes**

